# Even if suboptimal, novelty drives human exploration

**DOI:** 10.1101/2022.07.05.498835

**Authors:** Alireza Modirshanechi, Wei-Hsiang Lin, He A. Xu, Michael H. Herzog, Wulfram Gerstner

**Affiliations:** Brain-Mind Institute, School of Life Sciences, EPFL, Lausanne, Switzerland; School of Computer and Communication Sciences, EPFL, Lausanne, Switzerland; Helmholtz Munich, Munich, Germany; Max Planck Institute for Biological Cybernetics, Tübingen, Germany

## Abstract

Humans successfully explore their environment to find ‘extrinsic’ rewards, even when exploration requires several intermediate *reward-free* decisions. It has been hypothesized that ‘intrinsic’ rewards such as novelty guide this reward-free exploration. However, different intrinsic rewards lead to different exploration strategies, some prone to suboptimal attraction to irrelevant stochastic stimuli, sometimes called the ‘noisy TV problem.’ Here, we ask whether humans show a similar attraction to reward-free stochasticity and, if so, which type of intrinsic reward guides their exploration. We design a multi-step decision-making paradigm where human participants search for rewarding states in an environment with a highly stochastic but reward-free sub-region. We show that (i) participants persistently explore the stochastic sub-region and (ii) their decisions are best explained by algorithms driven by novelty but not by ‘optimal’ algorithms driven by information gain. Our results suggest that humans use suboptimal but computationally cheap strategies for exploration in complex environments.

## Introduction

Humans frequently search for more valuable rewards (e.g., more nutritious foods or better-paid jobs) than those currently available^1–3^. However, the computational and algorithmic nature of this exploratory behavior has remained highly debated^4–6^. State-of-the-art models of human exploration use Intrinsically Motivated Reinforcement Learning (RL) algorithms^7–10^ that, initially inspired by research in psychology^11,12^, have been designed to solve complex machine learning tasks with sparse ‘extrinsic’ rewards^13–19^. These algorithms use internally generated signals like ‘novelty,’ ‘surprise,’ or ‘information gain’ as ‘intrinsic’ rewards to guide exploratory action choices^11^. However, different intrinsic rewards result in different exploration strategies^20,21^. An unresolved yet crucial puzzle in neuroscience and psychology is identifying the type of intrinsic reward that drives exploration in humans^9,10^.

Resolving this puzzle primarily requires advances in experimental design. Specifically, experimental studies of human exploration have been mainly limited to simplistic experimental paradigms where a single action (or at most a pair of actions) is sufficient for reaching an extrinsic reward^22–28^or information^29–33^. These tasks are principally different from exploration in the real world where reaching a ‘goal’ requires several intermediate actions with no explicit progress feedback^9^. This has recently led to major concerns about the reliability and relevance of these tasks for characterizing human exploratory behavior^34–36^. Studying exploration in multi-step tasks^37,38^is hence pivotal for understanding and modeling human exploration^9,39,40^.

Compared to traditional experimental paradigms with homogeneously distributed stochasticity^41,42^, multi-step environments with a localized stochastic component have an important advantage since they enable the dissociation of exploration strategies based on different intrinsic rewards. Machine learning research has shown that intrinsically motivated RL agents are prone to distraction by stochasticity, i.e., they are attracted to novel, surprising, or just noisy states independently of whether or not these states are rewarding^43^ (the so-called ‘noisy TV’ problem^20,21^). However, the extent of this distraction varies between algorithms and depends on the type of intrinsic reward^44–48^. Artificial RL agents seeking *information gain* eventually lose their interest in stochasticity when exploration yields no further information^20,21^; in contrast, RL agents seeking *surprise* or *novelty* exhibit a persistent attraction by stochasticity^20,21^.

Here, we ask (i) whether humans are distracted in the same situations as intrinsically motivated RL agents and, if so, (ii) whether this distraction vanishes (similar to seeking information gain) or persists (similar to seeking surprise or novelty) over time.

## Results

We designed an experimental paradigm that dissociates different exploration strategies in an environment with 58 states plus three goal states (Fig. 1A-B). Three actions were available in each non-goal state, and agents could move from one state to another by choosing these actions (arrows in Fig. 1A-B). We use the term ‘agents’ to refer to either human participants or agents simulated by RL algorithms. In the human experiments, states were represented by images on a computer screen and actions by three disks below each image (Fig. 1C); for RL agents, both states and actions were abstract entities (i.e., we considered RL in a tabular setting^49^). The assignment of images to states and disks to actions was random but fixed throughout the experiment (Fig. 1C2). Agents were informed that there were three different goal states in the environment (*G*^*^, *G*_1_, or *G*_2_ in Fig. 1A) and that their task was to find a goal state 5 times; see Methods for how this information was incorporated in the RL algorithms. Neither human participants nor RL agents were aware of the total *number* of states or the *structure* of the environment (i.e., how states were connected).

**Figure 1:**
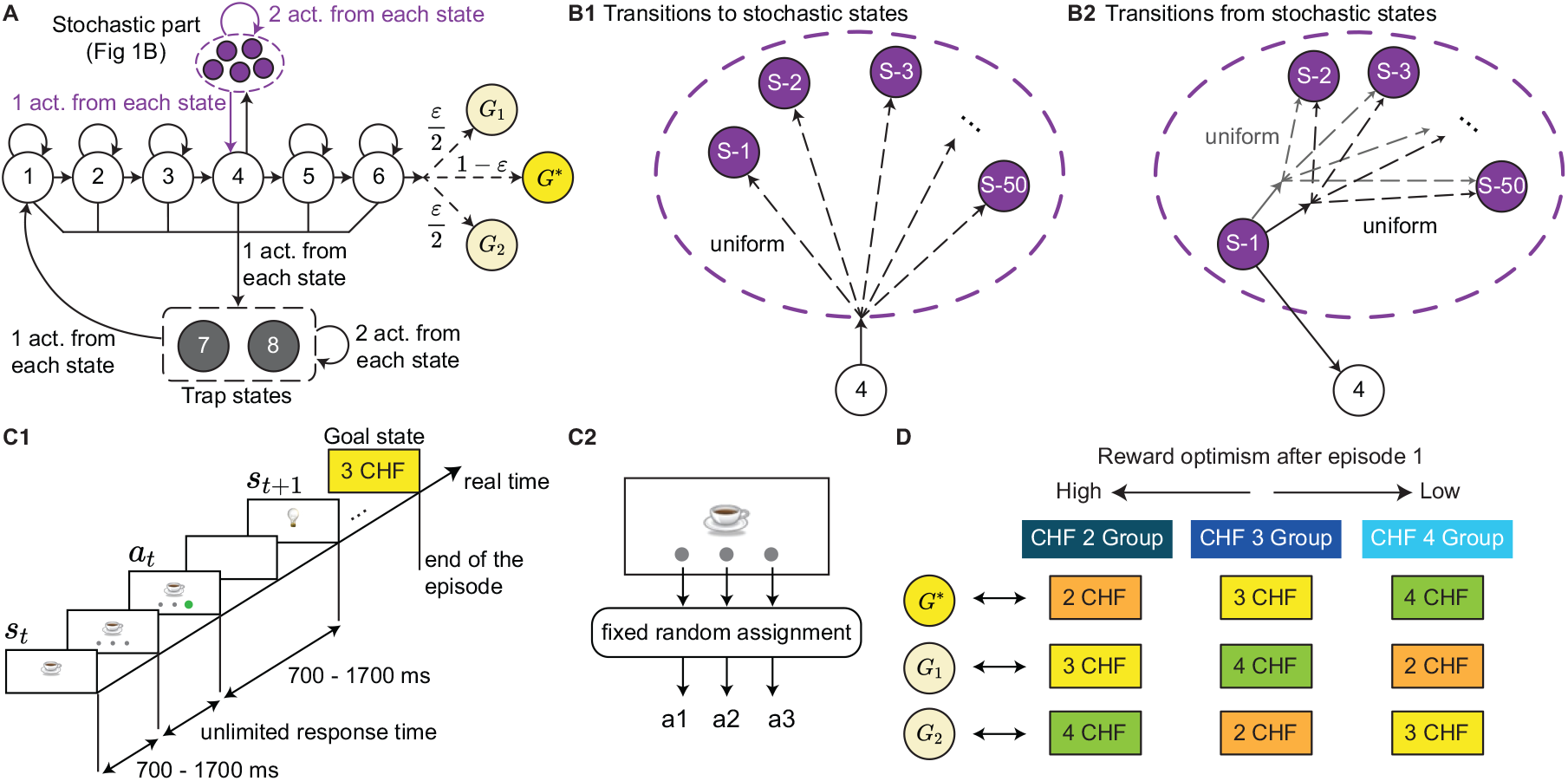
Experimental paradigm. **A**.Structure of the environment; only 5 out of the 50 stochastic states are shown (dashed oval; see B). Each circle represents a state and each solid arrow an action. All actions except those to the stochastic part or to the goal states are deterministic. Dashed arrows indicate random transitions; values (e.g., 1 − *ε*) show the probabilities of each transition. We chose *ε* ≪ 1 (see Methods). **B**. Zoom on stochastic transitions between states S-1 to S-50 inside the dashed oval. **B1**. In state 4, one action takes agents randomly (with uniform distribution) to one of the stochastic states. **B2**. In each stochastic state (e.g., state S-1 in the figure), one action (always the same) takes agents back to state 4 and two actions to another randomly chosen stochastic state. **C**. Timeline of one episode in human experiments (C1). The states were represented by images on a computer screen and actions by disks below each image. The assignment of images to states and disks to actions was random but fixed throughout the experiment (C2). An episode ended when a goal image (i.e., ‘3 CHF’ image in this example) was found. **D**. Human participants were informed that there were three goal states in the environment and that these goal states had different monetary values of 2 Swiss Franc (CHF), 3 CHF, and 4 CHF. For each participant, these monetary reward values were randomly assigned to different goal locations (i.e., *G*^*^, *G*_1_, and *G*_2_ in A) at the beginning of the experiment (without informing them); the assignment was fixed throughout the experiment. Hence, *G*^*^ had a different value for different participants, resulting in three groups of participants with different levels of reward optimism during episodes 2-5 (i.e., after finding *G*^*^ for the first time). See Methods.

The 58 states of the environment were classified into three groups: Progressing states (1 to 6 in Fig. 1A), trap states (7 and 8 in Fig. 1A), and stochastic states (S-1 to S-50 in Fig. 1B, shown as a dashed oval in Fig. 1A). In each progressing state, one action (‘progressing’ action) brought agents one step closer to the goals, while another (‘bad’ action) brought them to one of the trap states. The third action in states 1-3 and 5-6 was a ‘self-looping’ action that made agents stay in the same state. Except for the progressing action in state 6, all these actions were deterministic, meaning that they always led to the same next state. The progressing action in state 6 was *almost* deterministic: It took participants to the ‘likely’ goal state *G*^*^ with a probability of 1 − *ε* and to the ‘unlikely’ goal states *G*_1_ and *G*_2_ with equal probabilities of 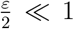. In state 4, instead of a self-looping action, there was a ‘stochastic’ action that took agents to a randomly chosen (with equal probability) stochastic state (Fig. 1B1). In each stochastic state, one fixed action (e.g., the left disk) reliably took agents back to state 4, and two stochastic actions took them to *another* randomly chosen stochastic state (Fig. 1B2). In each trap state, all three actions were deterministic: Two actions brought agents to either the same or the other trap state and one action to state 1.

The stochastic part of the environment – which mimics the main features of a ‘noisy TV’^43^ – is the crucial difference to existing paradigms^37,38,50,51^. Without the stochastic part, *all* types of intrinsic reward would help agents avoid the trap states and find the goal^37^. Hence, intrinsic rewards would help exploration before and not harm exploitation after finding a goal. However, the stochastic part dissociates exploratory behaviors driven by different intrinsic rewards; we elaborate on these differences in later sections (see ref.^20^ and Supplementary Materials).

### Reward optimism as an incentive to explore

We recruited 63 human participants and instructed them to perform our task for 5 episodes: Each episode started by initializing participants at state 1 or 2 and ended when they found any one of the 3 goal states (i.e., *G*^*^, *G*_1_, and *G*_2_). However, we chose a small enough *ε* (Fig. 1A) to safely assume that all participants would visit only *G*^*^ while being aware that *G*_1_ and *G*_2_ existed.

To further motivate exploration, we informed human participants that there were three different possible reward states corresponding to values of 2 Swiss Franc (CHF), 3 CHF, and 4 CHF, represented by three different images (see Methods for details and incorporating this information in the RL algorithms). At the beginning of the experiment, we randomly assigned the three different reward values to the goal states *G*^*^, *G*_1_, and *G*_2_, separately for each participant (without informing them), and kept the assignment fixed throughout the experiment (Fig. 1D). Following this random assignment, and after excluding 6 participants from further analyses (see Methods for criteria), *G*^*^ held different reward values across participants: 21 of 57 participants were assigned to environments with 2 CHF reward value for *G*^*^, 19 participants to environments with 3 CHF reward value for *G*^*^, and 17 participants to environments with 4 CHF reward value for *G*^*^. In the following, we refer to each group by their reward value of *G*^*^, e.g., the 3 CHF group is the group of human participants who had a reward value of 3 CHF for *G*^*^ (Fig. 1D).

The resulting three groups of human participants were characterized by three different levels of ‘reward optimism’ in episodes 2-5, where we define reward optimism as the expectancy of finding a goal of higher value than the one already discovered (Fig. 1D); we note that reward optimism in our experiment is closely linked to but independent of general optimism in psychology ^52^. Hence, even though all participants had received the same instructions, the 4 CHF group did not have any monetary incentive to explore further in episodes 2-5, whereas the 2 CHF group had a high monetary incentive to explore and find a higher reward in episodes 2-5. Therefore, we expected participants in the 2 CHF group to continue searching for more valuable goals in episodes 2-5. In the next sections, we aim to identify the dominant drive of this search behavior.

### Human participants persistently explore the stochastic part

We first studied the behavior of human participants without explicit computational modeling. During the 1st episode, all three groups of participants (i.e., 2 CHF, 3 CHF, and 4 CHF) had to explore the environment until they found the goal state *G*^*^ for the first time. Hence, their actions were solely exploratory. Importantly, they received no intermediate reward or progress feed-back throughout this exploration. Nevertheless, the participants learned to avoid the trap states (Fig. 2A, left) and were attracted to exploring the stochastic part of the environment (Fig. 2A, right). This suggests that participants used a guided exploration strategy (as opposed to a random exploration strategy).

**Figure 2:**
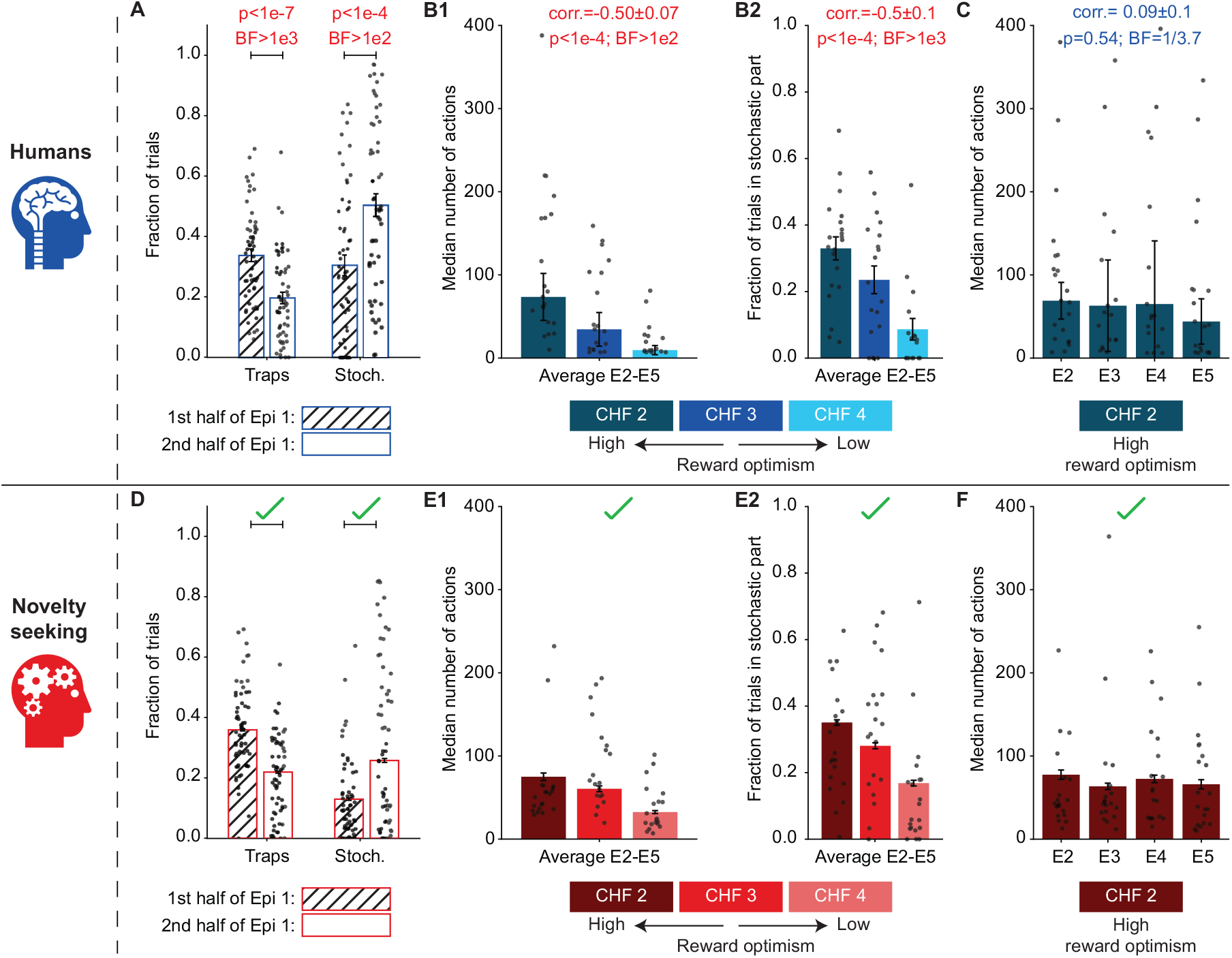
Human participants persistently explore the stochastic part. **A**.Participants spent less time in the trap states (one-sample t-test; *t* = −6.35; 95%CI = (−0.186, −0.097); DF = 56) and more time in the stochastic part (*t* = 4.25; 95%CI = (0.073, 0.203); DF = 56) during the 2nd half of episode 1 than during the 1st half. Error bars show the standard error of the mean (SEMean). **B**. Search duration in episodes 2-5. **B1**. Median number of actions over episodes 2-5 for the three different groups: 2 CHF (dark), 3 CHF (medium), and 4 CHF (light). Error bars show the standard error of the median (SEMed; evaluated by bootstrapping). The Pearson correlation between the search duration and the goal value is negative (correlation test; *t* = −4.2; 95%Confidence Interval (CI) = (−0.67, −0.27); Degree of Freedom (DF) = 55; Methods). **B2**. Average fraction of trials spent in the stochastic part of the environment during episodes 2-5. The Pearson correlation between the fraction of trials spent in the stochastic part and the goal value is negative (correlation test; *t* = −4.7; 95%CI = (−0.70, −0.32); DF = 55; Methods). Error bars show the SEMean. **C**. Median number of actions in episodes 2-5 for the 2 CHF group. A Bayes Factor (BF) of 1/3.7 in favor of the null hypothesis ^53^ suggests a zero Pearson correlation between the search duration and the episode number (one-sample t-test on individual correlations; *t* = 0.63; 95%CI = (−0.20, 0.37); DF = 20). Error bars show the SEMed. **D-F**. Posterior Predictive Check (PPC): Simulating novelty-seeking RL in our experimental paradigm successfully replicates the main qualitative patterns of the summary statistics of the action choices of human participants (see Fig. 4C for the quantification over 44 summary statistics). Panels D-F correspond to panels A-C, respectively, and illustrate the same summary statistics but for 1500 simulated novelty-seeking agents. Single dots in all panels show the data of individual human participants (A-C) or a subset (20 per group) of simulated participants (D-F). Red p-values in A-C: Significant effects with False Discovery Rate controlled at 0.05 ^54^ (see Methods). Red BFs in A-C: Significant evidence in favor of the alternative hypothesis (BF≥ 3). Blue BFs in A-C: Significant evidence in favor of the null hypothesis (BF≤ 1*/*3).

After finding the goal *G*^*^ for the 1st time (i.e., at the beginning of episode 2), each participant had effectively two options: (i) attempt to return to the discovered goal state *G*^*^ (exploitation) or (ii) search for the other goal states *G*_1_ and *G*_2_ (exploration). We quantified the extent of the exploratory behavior during episodes 2-5 by the search duration (i.e., the number of actions taken before returning to the discovered goal state; Fig. 2B1) and the fraction of trials spent in the stochastic part (Fig. 2B2). Both of these quantities were negatively correlated with the reward value of *G*^*^, e.g., the 2 CHF group had a longer search duration and spent more time in the stochastic part than the other two groups. Nevertheless, we still found a non-negligible exploration of the stochastic part by some participants in the 4 CHF group (Fig. 2B2, light blue), even though they had already found the goal state with the highest reward value. These observations (i) support the hypothesis that a higher degree of reward optimism leads to higher exploration in human participants and (ii) imply that human exploratory behavior is guided towards the stochastic part of the environment, even when there is no monetary incentive for exploration (see next section).

The behavior of the 2 CHF group is particularly interesting since, by design, they were the most optimistic group about finding higher rewards. The 2 CHF group exhibited a constant search duration over episodes 2-5 (zero correlation between the search duration and episode index confirmed by Bayesian hypothesis testing^53^; Fig. 2C). This implies that they persistently explored the stochastic part, even though it would have been theoretically possible to infer the structure of the environment and decrease exploration over time – as shown by ‘optimal’ agents seeking information gain (see ref.^20^ for a review and Supplementary Materials for simulations). Collectively, these results show that human exploration is not random but is also not theoretically optimal.

### Human participants successfully learned the environment’s structure

Thus far, we have shown that human participants exhibited a persistent attraction to the stochastic part in episodes 2-5, which is theoretically suboptimal. However, an implicit premise of our conclusion is that participants had learned the environment’s structure well enough to know how to return to *G*^*^ in episodes 2-5. To test this premise, we next analyzed whether participants could reconstruct the environment’s structure at the end of the experiment (Fig. 3). After finishing the experiment, participants were asked to reconstruct a map of the environment by connecting the images of different states (Fig. 3A; Methods). All three groups of participants achieved an above-chance reconstruction score (Fig. 3B; Methods), and a large majority of participants reconstructed the complete path from the trap states to state 6 (Fig. 3A). This implies that, by the end of the experiment, participants had built an explicit mental path for reaching the goal state *G*^*^.

**Figure 3:**
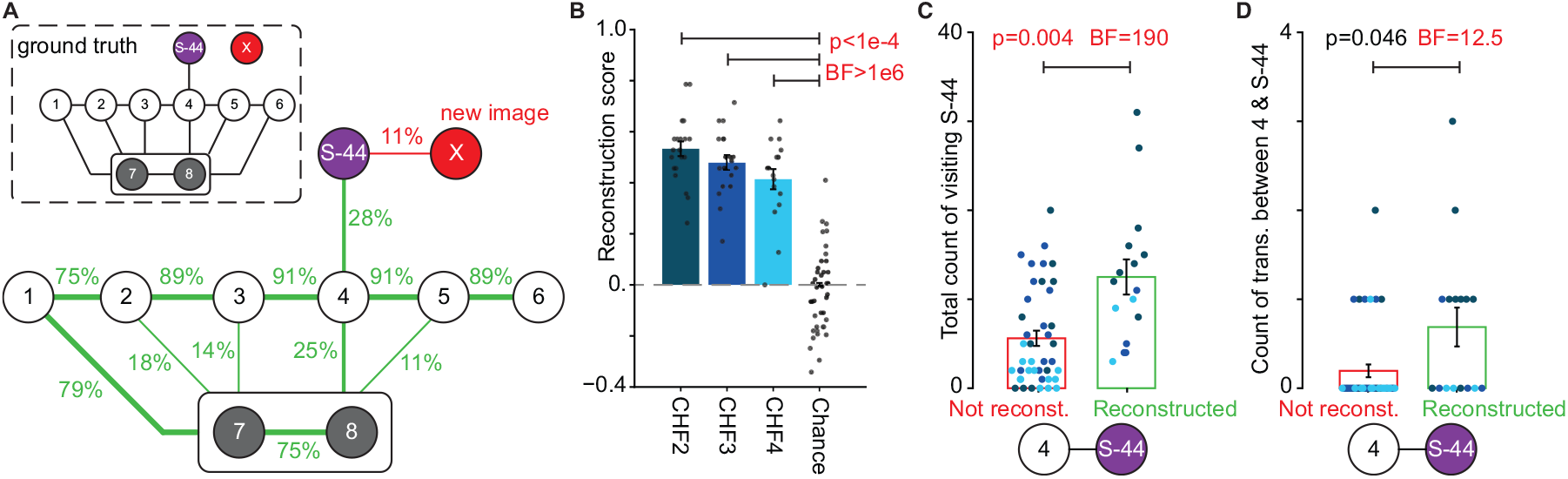
Human participants successfully reconstructed the environment’s underlying structure. At the end of the experiment, participants were presented with images of progressing states (1-6), trap states (7-8), one stochastic state (S-44), and a new image (X) that did not belong to the environment. The images were presented at once and in a pseudo-random spatial arrangement. Participants were asked to draw the experienced transitions between images (Methods). **A**. Average reconstruction of the environment’s structure. The reconstruction rate beside each link denotes the fraction of participants who drew that link. We only visualized links with a reconstruction rate higher than 10%. Inset: Ground truth. **B**. Reconstruction scores quantify the accuracy of participants’ reconstruction and take values between -1 (reconstructing only non-existing links) and +1 (perfect reconstruction; Methods). Random drawing yields on average a 0 reconstruction score (Chance). We observed a significantly above-chance reconstruction rate for participants in 2 CHF (one-sample t-test; *t* = 18.5; 95%CI = (0.47, 0.59); DF = 20), 3 CHF (*t* = 16.1; 95%CI = (0.42, 0.54); DF = 18), and 4 CHF groups (*t* = 10.5; 95%CI = (0.33, 0.50); DF = 17). **C-D**. Participants who reconstructed the link between state 4 and the stochastic state S-44 had visited S-44 significantly more often than those who did not (C; unequal variances t-test; *t* = 3.20; 95%CI = (2.4, 11.4); DF = 20.9); they had also experienced the transitions between states 4 and S-44 almost significantly more than those who did not (D; unequal variances t-test; *t* = 2.14; 95%CI = (0.01, 0.97); DF = 18.3). Red p-values in B-D: Significant effects with False Discovery Rate controlled at 0.05 ^54^ (see Methods). Red BFs in B-D: Significant evidence in favor of the alternative hypothesis (BF ≥ 3). Error bars in B-C: SEMean. Single dots in B-D: Data of individual participants (color-coded based on their reward group in C-D); for random drawing in B (Chance), we showed only 40 out of 1000 samples.

The images presented to participants also included one of the stochastic states (S-44) and a new image (X) that did not belong to the 58 states of the environment. Almost one-third of the participants successfully reconstructed the link between state 4 and the stochastic state, while *no* participants reconstructed a link between state 4 and the new image X (Fig. 3A). Importantly, while reconstructing the link between states 4 and S-44 indicates that the participant had learned the transition from state 4 to some stochastic states, not reconstructing this link can be due to reasons other than lack of understanding of the environment’s structure. For example, some participants might have ignored this link because they thought it was unimportant as it was not on the path to rewards, because they could not remember this very specific stochastic state, or because they never experienced a transition between state 4 and S-44. In fact, we observed that participants who reconstructed the link between states 4 and S-44 had visited state S-44 more frequently than those who did not (Fig. 3C). Strikingly, half of the participants who reconstructed the link had never directly experienced this specific transition (Fig. 3D). This indicates that these participants had learned the structure so thoroughly that they could generalize and reconstruct a link they had never directly encountered.

Overall, these results provide direct evidence that human participants were able to reconstruct a step-by-step map of the environment – despite the unprecedented complexity of the environment compared to other behavioral RL paradigms^42,50^. Hence, these results complement recent findings on human graph learning^55–57^ and, most importantly, imply that participants’ theoretically suboptimal exploration strategy is not an obvious consequence of poor graph learning.

### Computational modeling of human exploration

To uncover the algorithmic form of human exploration, we modeled human participants by intrinsically motivated RL agents who move in an environment with an unknown number of states by seeking extrinsic and intrinsic rewards (Fig. 4A). In this framework, intrinsic rewards are given to agents internally, whenever they encounter a ‘novel,’ ‘surprising,’ or ‘informative’ state. In contrast, extrinsic rewards are received only at the three goal states (see Methods for details). Specifically, at each time *t*, an agent observes state *s*_*t*_, evaluates an intrinsic reward value *r*_int,*t*_ (e.g., the novelty of state *s*_*t*_), and evaluates an extrinsic reward value *r*_ext,*t*_ (which is zero except at the goal states). Intrinsic and extrinsic reward values are then passed to two parallel, but separate, RL systems, each working with a single reward signal. Independently of each other, the two RL systems use a hybrid algorithm^37,50,58,59^combining model-based planning^60,61^and model-free habit-formation^62^ to learn a policy *π*_ext,*t*_ that maximizes future extrinsic rewards and a policy *π*_int,*t*_ that maximizes future intrinsic rewards^20,37^, respectively. The two policies are combined into a final policy *π*_*t*_ for taking the next action *a*_*t*_. The degree of exploration is high if *π*_int,*t*_ dominates *π*_ext,*t*_ during action selection. We assumed that ‘reward optimism’ influences the relative influence of *π*_int,*t*_ and *π*_ext,*t*_ on the final policy *π*_*t*_ and, as a result, the extent of exploration (Methods).

**Figure 4:**
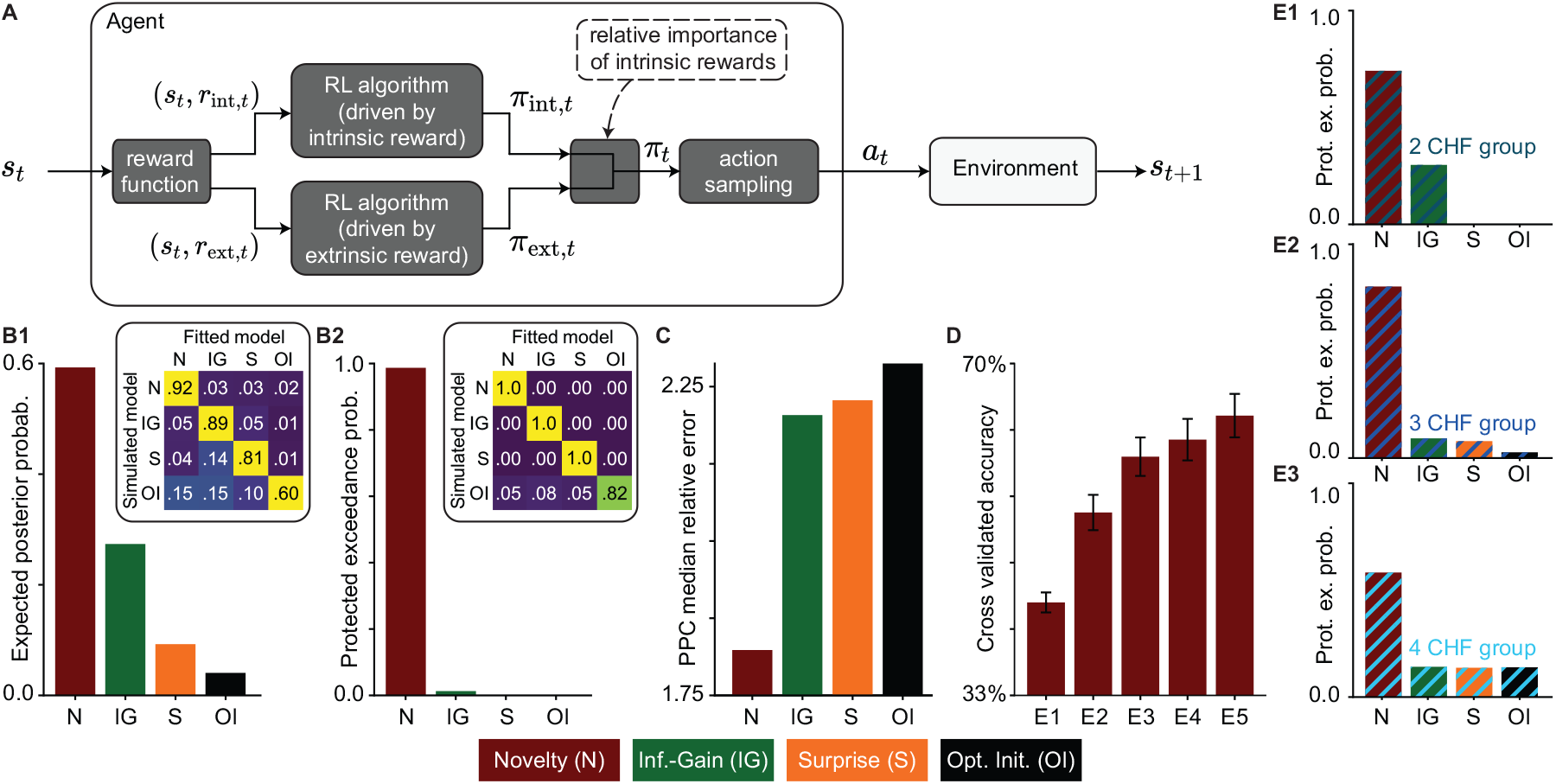
Novelty-seeking is the most accurate model of human behavior. **A**.Block diagram of the intrinsically motivated RL algorithm for modeling human behavior. Given the state *s*_*t*_ at time *t*, the intrinsic reward *r*_int,*t*_ (e.g., novelty) and the extrinsic reward *r*_ext,*t*_ (i.e., the monetary reward value of *s*_*t*_) are evaluated by a reward function and passed to two identical (except for the reward signals) parallel RL algorithms. The two algorithms compute two policies, one for seeking intrinsic reward *π*_int,*t*_ and one for seeking extrinsic reward *π*_ext,*t*_. The two policies are then weighted according to the relative importance of the intrinsic reward and are combined to make a final policy *π*_*t*_. The next action *a*_*t*_ is selected by sampling from *π*_*t*_. See Methods for details. **B**. Bayesian model comparison: Human participants’ action choices are best explained by novelty-seeking (N) compared to seeking information gain (IG), seeking surprise (S), or exploration based on optimistic initialization without intrinsic rewards (OI). **B1**. The expected posterior probability quantifies the proportion of participants whose behavior is best explained by each algorithm ^66^ (regarding cross-validated log-likelihoods; Methods). **B2**. Protected Exceedance Probability ^67^ quantifies the probability of each model being more frequent than the others among participants. Insets show confusion matrices from the model recovery ^68^ (see Methods); we could always recover the model that had generated the data, using almost the same number of simulated participants (60) as human participants (57). **C**. Model-comparison based on Posterior Predictive Checks (PPC): Median relative error (i.e., absolute difference divided by the SE) of each algorithm in replicating 44 group-level summary statistics of the action choices of human participants (e.g., fractions of trials spent in the stochastic part in Fig. 2A; see Methods for the full list). Novelty-seeking most accurately replicates human data. **D**. Cross-validated accuracy rate of novelty-seeking in predicting individual actions of human participants. The chance level is 33%. Error bars show the SEMean. Novelty-seeking allows above-chance prediction of each participant’s actions. **E**. Protected Exceedance Probability (as in B2) for participants in the 2 CHF (E1), 3 CHF (E3), and 4 CHF (E4) groups. Novelty-seeking is the most frequent model of behavior across and within groups.

We formulated three different hypotheses for human exploration in the form of three types of in- trinsic rewards *r*_int,*t*_; all three are representative examples of classes of intrinsic rewards in machine learning^20,21^: (i) novelty^13,14,37^, (ii) information gain^17,19,63,64^, and (iii) surprise^15,43,65^. Novelty quantifies how infrequent the state *s*_*t*_ has been until time *t*; thus, exploration in novelty-seeking agents is guided toward the least visited states. Information gain quantifies how much the agent updates its belief about the structure of the environment upon observing the transition from the state-action pair (*s*_*t*−1_, *a*_*t*−1_) to state *s*_*t*_; thus, exploration in information-gain-seeking agents is guided toward states where the agents’ estimates of the transition probabilities are least certain. Surprise quantifies how unexpected it is to observe state *s*_*t*_ after taking action *a*_*t*−1_ at state *s*_*t*−1_; thus, exploration in surprise-seeking agents is guided toward states with the most stochastic actions. As a control, we also considered the hypothesis that no explicit intrinsic reward signal is needed to explain human exploratory actions. We formalized this hypothesis in the form of an algorithm that uses no intrinsic rewards but incorporates some exploration incentive into the reward-seeking policy *π*_ext,*t*_ (via optimistic initialization^49^; see Methods).

### Novelty is the dominant drive of human exploration

To test which algorithm best explains human behavior, we used three-fold cross-validation^69^: We fitted the parameters of our four algorithms (i.e., novelty-seeking, information-gain-seeking, surprise-seeking, and exploration via optimistic initialization) to the action choices of two-thirds of human participants by maximizing the likelihood of data given model parameters (Methods). We then quantified the predictive power of the fitted algorithms by computing the likelihood of data for the rest of the participants using the fitted parameters (Methods).

Given the cross-validated likelihood of different algorithms, we used Bayesian model comparison^41,67^to rank the models (Methods). We find that seeking novelty is by far the most probable model for the majority of human participants, followed by seeking information gain as the 2nd most probable model (Fig. 4B; model-recovery^68^ in inset). Repeating the model comparison separately for each group of participants yielded the same conclusion (Fig. 4E; despite the ∼70% decrease in the sample size). This result shows (i) that seeking novelty describes the behavior of human participants better than seeking information gain, seeking surprise, or exploration via optimistic initialization and (ii) that reward optimism mainly influences the *extent* of the exploration but does not have a strong influence on the exploration *strategy*.

To confirm the results of our model comparison, we simulated each of the four algorithms with their fitted parameters in our experimental paradigm, i.e., we performed Posterior Predictive Checks (PPC)^68,70^. We then compared 44 summary statistics of human action choices (e.g., the fractions of trials spent in the stochastic part as in Fig. 2A) with those of the simulated agents (see Methods for the complete list of summary statistics). Results of the PPC show that novelty-seeking is *quantitatively* the most accurate algorithm in reproducing data statistics (Fig. 4C and Supplementary Materials). Novelty-seeking also successfully reproduced all key *qualitative* behavioral patterns of human participants discussed above (compare Fig. 2A-C with Fig. 2D-F).

Finally, to further test the predictive power of novelty-seeking, we quantified its accuracy in predicting individual actions of human participants (i.e., given a participant’s actions until time *t*, we asked whether novelty-seeking could predict the participant’s action at *t* + 1; Methods). We found a more than 40% cross-validated accuracy rate in episode 1 (Fig. 4D; chance level: 33%). As the participants moved through the environment, their behavior became more predictable (i.e., it was determined more strongly by their experience throughout the experiment than by their life experience before the experiment): Hence, we observed an increase in the cross-validated accuracy rate for episodes 2-5, with a more than 60% accuracy rate in episode 5. Therefore, novelty-seeking enabled an above-chance prediction of each participant’s actions, even though it had no prior information about the participant.

Taken together, our results provide strong quantitative and qualitative evidence for novelty as the dominant drive of human exploration in our experiment.

## Discussion

We designed a novel experimental paradigm to study human goal-directed exploration in multistep stochastic environments with sparse rewards. We made three main observations: (i) Human participants who were optimistic about finding higher rewards than those already discovered were persistently attracted to the stochastic part; (ii) the extent of attraction to the stochastic part decreased by decreasing the participants’ level of optimism, but it did not vanish even when there was no prospect of finding better rewards than the one already discovered; and (iii) this exploratory behavior was explained better by seeking novelty than seeking information gain or surprise, even though seeking information gain is theoretically more robust in dealing with stochasticity.

These three observations are instrumental in addressing the long-standing question of how humans explore their environments^4–6^. Specifically, past experimental studies have shown that humans use a combination of random and directed exploration in 1-step or 2-step decision-making tasks (e.g., multi-armed bandits)^22–24,71–73^, while theoretical studies have proposed distinct motivational signals as potential drives of human directed exploratory actions^5,8,9,74,75^. However, despite significant advances in the past years^25–27,29–31,76–83^, it has remained highly debated which motivational signal explains human exploration best^9,10^. Importantly, the focus of existing studies on 1-step or 2-step decision-making tasks has raised questions about whether our current understanding of human exploration can be generalized to more complex and realistic situations ^9,34–36,39^.

To bridge between exploration in 1-step and multi-step tasks, we showed in an earlier study^37^ that novelty dominantly drives human exploration in complex but *deterministic* environments with sparse rewards, i.e., situations where novelty-seeking has empirically been shown to be an effective exploration strategy ^13,14^. Observations (i)-(iii) above provide further evidence for novelty as the dominant drive of human goal-directed exploration even in situations *when seeking novelty is not optimal*. Specifically, after episode 1, participants can reasonably assume that the task is solvable, i.e., if they have succeeded in finding the 2 CHF reward, then they should also be able to find the higher rewards. Hence, the fact that the participants in the 2 CHF group continue the search during episodes 2-5 is expected and economically rational, but our results show that they use a *suboptimal* novelty-based search strategy. Further experimental studies are needed to investigate the implications of our results for other types of human exploratory behavior. In particular, it is a priori unclear whether goal-directed exploration, as studied here, shares some drives and mechanisms with reward-free exploration strategies in, e.g., reactive orienting and passive viewing ^80,84^, navigation^85,86^, and non-instrumental decision-making tasks^29,32,33^.

Our results appear to contradict the long-lasting belief that humans are not prone to the ‘noisy TV’ problem^1,46,48^. It is important, however, to note that the stochasticity in our environment is different from passively watching a noisy, grey-flickering TV screen. Rather, the environment allows participants to take actions that are in spirit similar to exploring different TV channels, where each channel contains videos – similar to the recent realizations of ‘noisy TV’ in machine learning^43^. In this context, our experimental paradigm is a model experiment of recent social media where users spend hours on the ‘endless scrolling option’ to watch new videos^87,88^– despite the availability of alternative activities with ‘extrinsic’ rewards. This is analogous to the behavior of the 4 CHF participants who kept exploring the stochastic part despite knowing the path to the most rewarding goal state.

Accordingly, our results challenge the optimality of human exploration^11,83^. However, we note that, for computing novelty, an agent only needs to track the state frequencies over time and does not need any knowledge of the environment’s structure (Methods); hence computing novelty is computationally cheaper than computing information gain. This suggests that a potentially higher level of distraction by novelty in humans may be the price of spending less computational power. In other words, novelty-seeking in the presence of stochasticity may not be a globally optimal strategy for exploration but can be an optimal strategy given a set of prior assumptions and computational constraints, i.e., a ‘resource rational’ policy ^89–91^.

Finally, we note that notions of ‘novelty’, ‘surprise’, and ‘information gain’ as scientific terms often refer to different precise mathematical definitions^65,92^ – across a broad set of applications in neuroscience^37,93,94^, psychology^95–97^, and machine learning^20,21,48^. Our results in this paper are based on the specific mathematical formulations that we have chosen (Methods), but we expect our conclusions to be invariant to the precise choice of definitions as long as (i) novelty quantifies infrequency of *states* ^37^ as, for example, defined with density models in machine learning^13,14,98^; (ii) surprise quantifies mismatches between observations and agents’ expectations, where the expectations are made based on the previous *state-action* pair, including all measures of prediction surprise^65^ and typical measures of prediction error in machine learning^15,43^; and (iii) information gain quantifies improvements in the agents’ *world-model* and vanishes with the accumulation of experience, which includes Bayesian^93^ and Postdictive surprise^94^, measures of disagreement and progress-rate in machine learning^17–19,44,99^, and optimal exploration bonuses in RL theory^100,101^.

In conclusion, our results show (i) that human decision-making is influenced by an interplay of intrinsic and extrinsic rewards, controlled by reward optimism, and (ii) that novelty-seeking RL algorithms can successfully model this interplay in tasks where humans search for rewarding states.

## Methods

### Ethics statement

The data for human experiment were collected under CE 164*/*2014, and the protocol was approved by the ‘Commission cantonale d’éthique de la recherche sur l’être humain’. All participants were informed that they could quit the experiment at any time and signed a written informed consent. All procedures complied with the Declaration of Helsinki (except for pre-registration).

### Experimental procedure

63 participants joined the experiment. Data from 6 participants were removed (see below); thus, data from 57 participants (27 female, mean age 24.1 ± 4.1 years) were included in the analyses. All participants were naive to the purpose of the experiment and had normal or corrected-to-normal visual acuity. The experiment was scripted in MATLAB using the Psychophysics Toolbox ^102^.

Before starting the experiment, the participants were informed that they needed to find any of the 3 goal states 5 times. They were shown the 3 goal images and informed that each image had a different reward value of 2 CHF, 3 CHF, or 4 CHF. Specifically, they were given an example that ‘if you find the 2 CHF goal twice, 3 CHF goal once, and 4 CHF goal twice, then you will be paid 2 × 2 + 1 × 3 + 2 × 4 = 15 CHF’; see Informing RL agents of different goal states for how this information was incorporated into the RL algorithms. At each trial, participants were presented with an image (state) and three grey disks below the image (Fig. 1C). Clicking on a disk (action) led participants to a subsequent image, which was chosen based on the underlying graph of the environment in Fig. 1A-B (which was unknown to the participants). Participants clicked through the environment until they found one of the goal states, which finished an episode (Fig. 1C).

The assignment of images to states and disks to actions was random but kept fixed throughout the experiment and identical for all participants (Fig. 1C2). Exceptionally, we did not make the assignment for the actions in state 4 before the start of the experiment. Rather, for each participant, we assigned the disk that was chosen in the 1st encounter of state 4 to the stochastic action and the other two disks randomly to the bad and progressing actions, respectively (Fig. 1A). With this assignment, we ensured that all human participants would visit the stochastic part at least once during episode 1. The same protocol was used for simulated RL agents. Additionally, to ensure that participants would not get lost in the stochastic part, we used the same assignment for the ‘escape action’ in all stochastic states (i.e., the action that took participants from stochastic states to state 4 in Fig. 1B).

Before the start of the experiment, we randomly assigned the different goal images (corresponding to the three reward values) to different goal states *G*^*^, *G*_1_, and *G*_2_, separately for each participant (Fig. 1D). The image and, hence, the reward value were then kept fixed throughout the experiment. In other words, we randomly assigned different participants to different environments with the same structure but different assignments of reward values. We, therefore, ended up with 3 groups of participants: 23 in the 2 CHF group, 20 in the 3 CHF group, and 20 in the 4 CHF group (Fig. 1D). The probability of encountering a goal state other than *G*^*^ was controlled by the parameters *ε*. We considered *ε* to be around machine precision 10^−8^, so we have (1 − *ε*)^5×63^ ≈ 1 − 10^−5^ ≈ 1, meaning that all 63 participants would be taken almost surely to the goal state *G*^*^ in all 5 episodes. We note, however, that a participant could, in principle, observe any of the 3 goals if they could choose the progressing action at state 6 sufficiently many times.

Two participants (in the 2 CHF group) did not finish the experiment, and four participants (1 in the 3 CHF group and 3 in the 4 CHF group) took more than 3 times group-average number of actions in episodes 2-5 to finish the experiment. We considered this as a sign of being non-attentive and removed these 6 participants from further analyses.

At the end of the experiment, participants were given a paper with the pseudo-randomly placed images of progressing states (1-6), trap states (7-8), one stochastic state (S-44), as well as a new image (X) that did not belong to the 58 states of the environment. Participants were asked to ‘draw the transitions between images’ and were told they ‘can add anything [they] want.’ Some participants had not reported the directionality of transitions. Hence, we only analyzed how many participants had drawn a link between every pair of states, independently of the link’s direction (Fig. 3). To further simplify analyses, we did not dissociate between different trap states when counting the connections from non-trap states to the trap states. As a result, there were 1 + 9 × 8*/*2 = 37 possible links to draw (the extra 1 belongs to the connection between the two trap states), but there were only 13 links in the ground truth (Fig. 3A, inset). Accordingly, we defined the reconstruction score in Fig. 3 as the ratio of correctly reconstructed links (out of 13) minus the ratio of incorrectly reconstructed links (out of 24).

The correction for multiple hypotheses testing was done by controlling the False Discovery Rate at 0.05^54^ over all 10 null hypotheses that are presented in Fig. 2 and Fig. 3 (p-value threshold: 0.045). Using Bonferroni correction (with a family-wise error rate of 0.05, i.e., p-value threshold: 0.005) does not change our results. All Bayes Factors (abbreviated BF in the figures) were evaluated using the Schwartz approximation ^53^ to avoid any assumptions on the prior distribution.

### Computational modeling

We used ideas from non-parametric Bayesian inference^103^ to design an intrinsically motivated RL algorithm for environments where the total number of states is unknown. We present the final results here and present the derivations and pseudo-code in Supplementary Materials.

We indicate the sequence of actions and states until time *t* by *s*_1:*t*_ and *a*_1:*t*_, respectively, and define the **set of all known states** at time *t* as

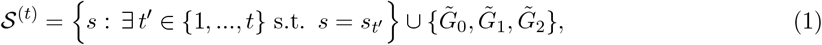

Where 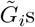 represent our three different goal states 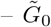 corresponds to the 2 CHF goal, 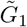 to the 3 CHF goal, and 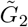 to the 4 CHF goal. Note that 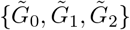 represents the images of the goal states and not their locations *G*^*^, *G*_1_, and *G*_2_ and that the assignment of images to locations is unknown to the model. Hence, starting with *t* = 0, the algorithm incorporates information about the existence of multiple goal states in the environment. In a more general setting, 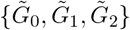 should be replaced by the set of all states whose images were shown to participants before the experiment. After a transition to state *s*_*t*+1_ = *s*^′^ resulting from taking action *a*_*t*_ = *a* ∈ {left, middle, right} (i.e., representing disk positions in Fig. 1C) at state *s*_*t*_ = *s*, the reward functions *R*_ext_ and *R*_int,*t*_ evaluate the reward values *r*_ext,*t*+1_ and *r*_int,*t*+1_. We define the **extrinsic reward function** *R*_ext_ as

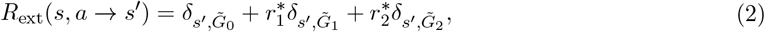

where δ is the Kronecker delta function, and we assume (without loss of generality) a subjective extrinsic reward value of 1 for 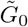 (2 CHF goal) and subjective extrinsic reward values of 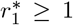 and 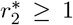 for 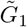 and 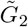, respectively. The prior information of human participants about the difference in the monetary reward values of different goal states can be modeled in simulated RL agents by varying 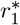 and 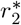 (resulting in the exploratory component of reward-seeking in optimistic initialization; see ‘Informing RL agents of different goal states‘). We discuss *R*_int,*t*_ in Alternative algorithms.

As a general choice for the RL algorithm in Fig. 4A, we consider a hybrid of model-based and model-free policy ^37,50,59,62^. The **model-free (MF) component** uses the sequence of states *s*_1:*t*_, actions *a*_1:*t*_, extrinsic rewards *r*_ext,1:*t*_, and intrinsic rewards *r*_int,1:*t*_ (in the two parallel branches in Fig. 4A) and estimates the extrinsic and intrinsic *Q*-values 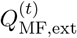 and 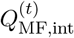, respectively. Traditionally, MF algorithms do not need knowledge of the total number of states ^49^; thus, the MF component of our algorithm remains similar to that of previous studies ^37,104^: At the beginning of episode 1, *Q*-values are initialized at 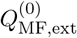 and 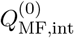. Then, the estimates are updated recursively after each new observation. After the transition (*s*_*t*_, *a*_*t*_) → *s*_*t*+1_, the agent computes extrinsic and intrinsic reward prediction errors *RPE*_ext,*t*+1_ and *RPE*_int,*t*+1_, respectively:

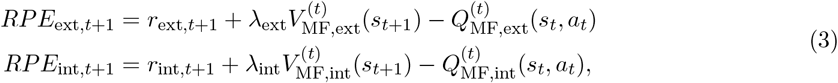

where *λ*_ext_ and *λ*_int_ ∈ [0, 1) are the discount factors for extrinsic and intrinsic reward seeking, respectively, and 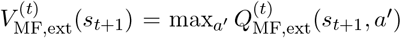 and 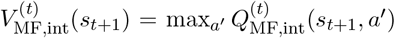 are the extrinsic and intrinsic *V* -values ^49^ of the state *s*_*t*+1_, respectively. We use two separate eligibility traces^49,104^for the update of *Q*-values, one for extrinsic reward *e*_ext,*t*_ and one for intrinsic reward *e*_int,*t*_, both initialized at zero at the beginning of each episode. The update rules for the eligibility traces after taking action *a*_*t*_ at state *s*_*t*_ is

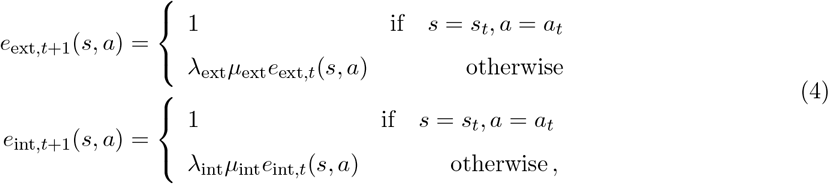

where *λ*_ext_ and *λ*_int_ are the discount factors defined above, and *µ*_ext_ and *µ*_int_ ∈ [0, 1] are the decay factors of the eligibility traces for the extrinsic and intrinsic rewards, respectively. The update rule is then 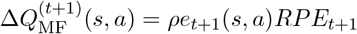, where *e*_*t*+1_ is the eligibility trace (i.e., either *e*_ext,*t*+1_ or *e*_int,*t*+1_), *RPE*_*t*+1_ is the reward prediction error (i.e., either *RPE*_ext,*t*+1_ or *RPE*_int,*t*+1_), and *ρ* ∈ [0, 1) is the learning rate.

The **model-based (MB) component** builds a world-model that summarizes the structure of the environment by estimating the probability *p*^(*t*)^(*s*^′^|*s, a*) of the transition (*s, a*) → *s*^′^. To do so, an agent counts the transition (*s, a*) → *s*^′^ recursively and using a leaky integration^105,106^:

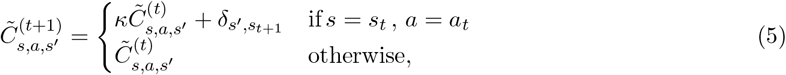

where δ is the Kronecker delta function, 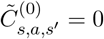, and *κ* ∈ [0, 1] is the leak parameter and accounts for imperfect memory during model-building in humans. If *κ* = 1, then 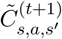 is the exact count of transition (*s, a*) → *s*^′^. For *κ <* 1, we refer to 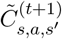 as a leaky count or pseudo-count. These leaky counts are used to estimate the transition probabilities

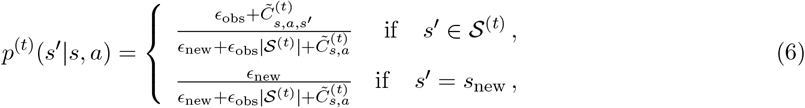

where 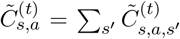 is the leaky count of taking action *a* at state *s, ϵ* _obs_ ∈ ℝ ^+^ is a free parameter for the prior probability of transition to a known state (i.e., states in S^(*t*)^), and *ϵ*_new_ ∈ ℝ ^+^ is a free parameter for the prior probability of transition to a new state (i.e., states not in S^(*t*)^) – see Supplementary Materials for derivations. Choosing *ϵ*_new_ = 0 is equivalent to assuming there is no unknown state in the environment, for which the estimate in Eq. 6 is reduced to the classic Bayesian estimate of transition probabilities in bounded discrete environments ^37,59^. The transition probabilities are then used in a novel variant of prioritized sweeping ^49,60^adapted to deal with an unknown number of states. The prioritized sweeping algorithm computes a pair of *Q*-values, i.e., 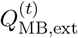 for extrinsic and 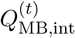 for intrinsic rewards, by solving the corresponding Bellman equations ^49^ with *T*_*P S*,ext_ and *T*_*PS*,int_ iterations, respectively. See Supplementary Material for details.

Finally, actions are chosen by **a softmax policy** ^49^: The probability of taking action *a* in state *s* at time *t* is

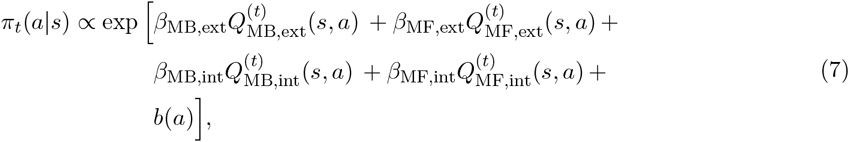

where *β*_MB,ext_ ∈ ℝ^+^, *β*_MF,ext_ ∈ ℝ^+^, *β*_MB,int_ ∈ ℝ^+^, and *β*_MF,int_ ∈ ℝ^+^ are free parameters (i.e., inverse temperature parameters of the softmax policy^49^) expressing the contribution of each *Q*-value to action-selection, and *b*(*a*) captures the general bias of the agent for taking the particular action *a* (e.g., left grey disk in Fig. 1C) independently of the state *s*. Without loss of generality, we assume *b*(left) = 0 and considered *b*(middle) ∈ ℝ and *b*(right) ∈ ℝ as free parameters. For Fig. 4A, we defined hybrid policies for each of the two branches as

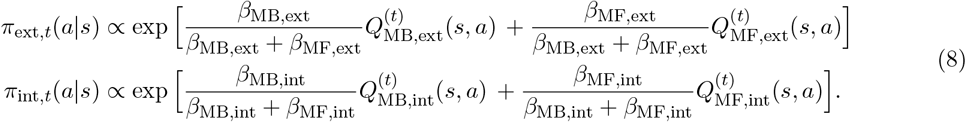

Hence the final policy is 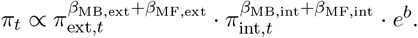.

In general, the contribution of seeking extrinsic reward and seeking intrinsic reward as well as the MB and MF branches to action selection depends on different factors, including time passed since the beginning of the experiment^51,62^, cognitive load ^107^, and whether the location of reward is known ^37^. Here, we make a simplistic assumption that these contributions (expressed as the 4 inverse temperature parameters) depend only on reward optimism:

- Episode 1: Before finding the goal state, we consider 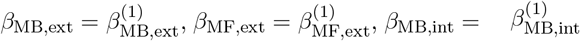, and 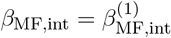 as four independent free parameters.
- Episodes 2-5: After finding the goal *G*^*^, we consider 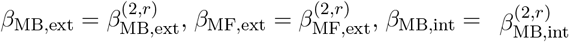, and 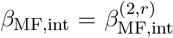, where *r* is either 2 CHF, 3 CHF, or 4CHF, resulting in 3 × 4 = 12 free parameters.

#### Summary of free parameters

The full algorithm has 14 main parameters (capturing initialization and learning dynamics)

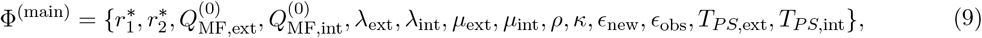

16 inverse temperature parameters (capturing the randomness in decision-making and the balance of seeking intrinsic versus extrinsic rewards)

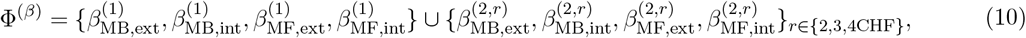

and 2 bias parameters

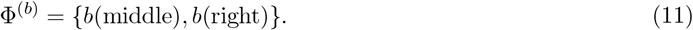

We denote the set of all parameters by

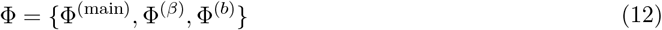

We note that *not* all these parameters were fitted for all algorithms (see Alternative algorithms).

### Informing RL agents of different goal states

Human participants were informed that the environment had different goal states with different monetary reward values. This information was intended to incentivize exploration after finding the likely goal state *G*^*^ at the end of episode 1. We used three mechanisms to incorporate this information into the RL algorithm described above (Computational modeling). Our main focus throughout the paper has been on the first mechanism: Reward optimism balances intrinsic rewards against extrinsic rewards (Fig. 4A). We formalized this idea by assigning different values to *β*_MB,ext_, *β*_MF,ext_, *β*_MB,int_, and *β*_MF,int_ (see Eq. 7) depending on the reward value of *G*^*^; this makes **the relative importance of intrinsic rewards** explicitly depend on the difference between the reward value of the discovered goal *r*_*G*_^*^ and the known reward values 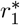 and 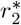 of the other goal states (Eq. 2).

The two other alternative mechanisms are the **model-based optimistic initialization** and **model-free optimistic initialization**. Exploration in the optimistic initialization algorithm in Fig. 4 is solely directed via these mechanisms (see Alternative algorithms). In this section, we discuss how these mechanisms balance exploration versus exploitation.

#### Model-based optimistic initialization

MB optimistic initialization is an explicit approach to model reward-optimism through designing the world-model. The MB branch finds the extrinsic *Q*-values 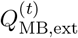 by solving the Bellman equations

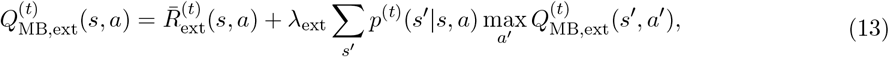

where *p*^(*t*)^(*s*^′^|*s, a*) is estimated transition probability in Eq. 6, and

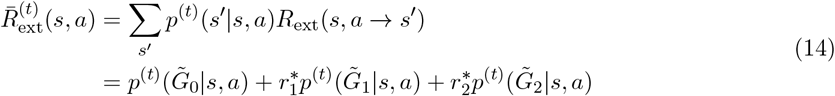

is the average immediate extrinsic reward expected to be collected by taking action *a* in state *s* (see Eq. 2). Hence, the knowledge of the existence of three different goal states with three different rewards has an explicit influence on the MB branch. For example, because no transitions to any of the goal states have been experienced during episode 1, we have

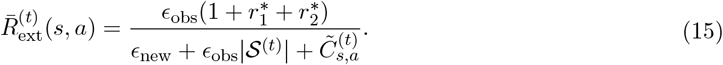

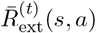 is closely linked to (approximately) Bayes-optimal exploration bonuses in the RL theory^100^ and has two important properties. First, 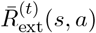 is an increasing function of *ϵ*_obs_. This implies that the expected reward of a transition during episode 1 increases by increasing the prior probability of transition to states in 𝒮^(*t*)^. This is a direct consequence of our Bayesian approach to estimating the world-model. Second, 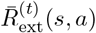 is a decreasing function of 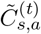. This implies that the expected reward of a state-action pair decreases by experience. Importantly, 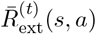 converges to 0 as 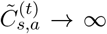, which makes a link between exploration driven by the MB optimistic initialization and exploration driven by information gain (see below).

During episodes 2-5, the exact theoretical analysis of the MB optimistic initialization is rather complex. However, using a few approximation steps for episode 2, we can find a condition for whether the MB extrinsic *Q*-values show a preference for exploring or leaving the stochastic part (Supplementary Materials). The condition involves a comparison between the discounted reward value of the discovered goal state 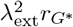 and an optimistic estimate of a reward-to-be-found 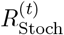 in the stochastic part that depends on 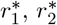, *λ*_ext_, *ϵ*_obs_, |𝒮^(*t*)^|, and the average pseudo-count 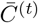 of state-action pairs in the stochastic part (Supplementary Materials). We can show that if 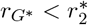, then increasing 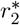 would eventually result in a preference for staying in the stochastic part: If the reward value of a goal state is much greater than the value of the discovered goal state, then the agent prefers to keep exploring the stochastic part. However, for any value of 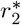 and *r*_*G*_^*^, increasing 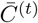 would eventually result in a preference for leaving the stochastic part and going towards the already discovered goal: Agents will eventually give up exploration after a sufficiently long and unsuccessful exploration phase. This is another qualitative link between exploration based on the MB optimistic initialization and exploration driven by information gain (see below).

#### Model-free optimistic initialization

Unlike the MB branch, the MF branch does not explicitly know about the existence of different goal states and their values. However, the initial value 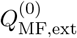 of the MF extrinsic *Q*-values quantifies an expectation of the reward values in the environment before any interaction with the environment. During episode 1, no extrinsic reward is received by the agent; hence, for a small enough learning rate *ρ* and an optimistic initialization 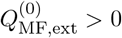, the extrinsic reward prediction errors are always negative (Eq. 3). As a result, 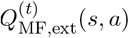 decreases as an agent keeps taking action *a* in state *s*, which motivates the agent to try new actions. This is a well-known mechanism for directed exploration in the machine learning community^49^. Similar to the MB optimistic initialization, the effect of the MF optimistic initialization fades out over time – which makes them both similar to exploration driven by information gain (see below).

During episodes 2-5, the exact theoretical analysis of the MF optimistic initialization is complex and dependent on an agent’s exact trajectory (because of the eligibility traces). However, whether the MF extrinsic *Q*-values show a preference for exploring or leaving the stochastic part essentially depends on the reward value of the discovered goal state *r*_*G*_^*^ and the initialization value 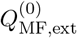. For example, if an agent, starting at *s*1, takes the perfect trajectory of *s*1 → *s*2 → *s*3 → *s*4 → *s*5 → *s*6 → *G*^*^ in episode 1, then, given a unit decay factor of the eligibility traces (i.e., *µ*_ext_ = 1), it is easy to see that, in the 1st visit of state 4 in episode 2, the agent prefers the stochastic/bad action over the progressing action if 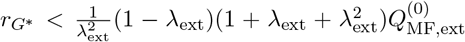. This implies that, even though the MF branch is not explicitly aware of different goal states and their reward values, it can still describe a type of reward optimism through the initialization of *Q*-values.

### Alternative algorithms

We considered four hypotheses for how humans explore the environment to search for the goal state (including most representative explorations strategies in RL ^9,20,21^): (i) seeking novelty, (ii) seeking information gain, (iii) seeking surprise, and (iv) exploration via optimistic initialization (i.e., no intrinsic rewards). We formalized the four hypotheses in our framework by using different types of the intrinsic reward function *R*_int,*t*_ that maps a transition (*s, a*) → *s*^′^ to an intrinsic reward value *r*_int,*t*+1_ = *R*_int,*t*_(*s*_*t*_, *a*_*t*_ → *s*_*t*+1_). In this section, we describe these algorithms.

#### 1. Novelty-seeking

For an agent seeking novelty (red in Fig. 4), we defined the intrinsic reward function as

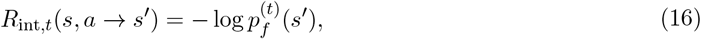

where 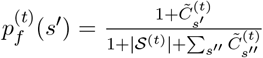 is the state frequency with 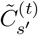 the pseudo-count of encounters of state *s*^′^ up to time *t* (similar to Eq. 5): 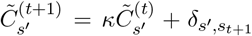 with 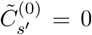. With this definition, that generalizes earlier works ^37^ to the case where the number of states is unknown, the least novel states are those that have been encountered most often (i.e., with the highest 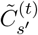). Moreover, novelty is at its highest value for the unobserved states as we have 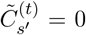 for any unobserved state *s*^′^ ∉ 𝒮^(*t*)^. Similar intrinsic rewards have been used in machine learning^13,14^.

To dissociate the effect of exploration by novelty-seeking from optimistic initialization in episode 1, we considered 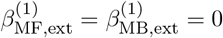 and 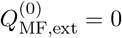. Moreover, we put *T*_*P S*,ext_ = *T*_*P S*,int_ = 100 (i.e., almost twice the total number of states) to decrease the number of parameters, based on the results of ref. ^37^ showing the negligible importance fitting this parameter. Hence, the novelty-seeking algorithm had a total of **27 parameters** (11 main parameters + 14 inverse temperature parameters + 2 biases).

#### 2. Information-gain-seeking

For an agent seeking information gain (green in Fig. 4), we defined the intrinsic reward function as

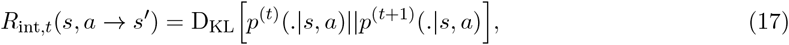

where D_KL_ is the Kullback-Leibler divergence^108^, and *p*^(*t*+1)^ is the updated world-model upon observing (*s, a*) → *s*^′^. The dots in Eq. 17 denote the dummy variable over which we integrate to evaluate the Kullback-Leibler divergence. Note that if *s*^′^ ∉ 𝒮^(*t*)^, then there are some technical problems in the naive computation of D_KL_ – since *p*^(*t*)^ and *p*^(*t*+1)^ have different supports. We dealt with these problems using a more fundamental definition of D_KL_ using the Radon–Nikodym derivative; see Supplementary Materials for derivations and see ref. ^63^ for an alternative heuristic solution. Note that the information gain in Eq. 17 has also been interpreted as a measure of surprise (called ‘Postdictive surprise’ ^94^), but it has a behavior radically different from that of the Shannon surprise introduced below for our surprise-seeking agents (Eq. 18) – see ref. ^65^ for an elaborate treatment of the topic. Importantly, the expected (integrated over *s*^′^) information gain corresponding to a state-action pair (*s, a*) converges to 0 as 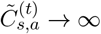 (see Supplementary Materials for the proof). Similar intrinsic rewards have been used in machine learning^17,44,48,63^.

Similarly to the case of novelty-seeking, we considered 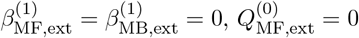, and *T*_*P S*,ext_ = *T*_*P S*,int_ = 100; hence, the algorithm seeking information gain also had a total of **27 parameters** (11 main parameters + 14 inverse temperature parameters + 2 biases).

#### 3. Surprise-seeking

For an agent seeking surprise (orange in Fig. 4), we defined the intrinsic reward function as the Shannon surprise (a.k.a. surprisal) ^65^

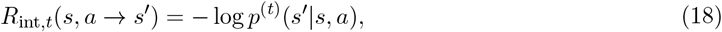

where *p*^(*t*)^(*s*^′^|*s, a*) is defined in Eq. 6. With this definition, the expected (integrated over *s*^′^) intrinsic reward of taking action *a* at state *s* is equal to the entropy of the distribution *p*^(*t*)^(*s*^′^|*s, a*) ^108^. If *ϵ*_new_ *< ϵ*_obs_, then the most surprising transitions are the ones to unobserved states. Similar intrinsic rewards have been used in machine learning^15,43^.

Similarly to the case of novelty-seeking, we considered 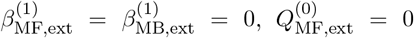, and *T*_*P S*,ext_ = *T*_*P S*,int_ = 100; hence, the surprise-seeking algorithm had also a total of **27 parameters** (11 main parameters + 14 inverse temperature parameters + 2 biases).

#### 4. Exploration by optimistic initialization (no intrinsic rewards)

As our last alternative algorithm (black in Fig. 4), we considered agents with no intrinsic reward:

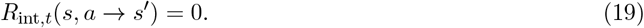

Exploratory actions of these agents are purely driven by MB and MF optimistic initialization described in Informing RL agents of different goal states. As a result, exploration based on optimistic initialization does not depend on any of the parameters that influence the intrinsically motivated part of the RL algorithm described above, ending up with a total of **21 parameters** (11 main parameters + 8 inverse temperature parameters + 2 biases) for the optimistic initialization (considering *T*_*P S*,ext_ = 100).

### Model-fitting and model-comparison

To compare different algorithms based on their explanatory power, we did a stratified 3-fold cross-validation ^69^: We grouped our 57 human participants into 3 disjoint sets, where all sets had almost the same number of participants from different reward groups (i.e., 2 CHF, 3 CHF, 4 CHF). For each fold *k* ∈ {1, 2, 3} of cross-validation, one set of participants was considered as testing set 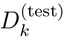 and the union of the other two as the training set 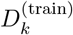.

Then, for each model *M* ∈ {novelty, inf-gain, surprise, opt. init.} and cross-validation fold *k* ∈ {1, 2, 3}, we fitted the model parameters Φ_*M*_ by maximizing likelihood of the training data given parameters:

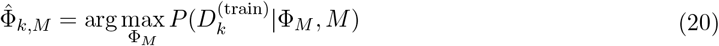

where 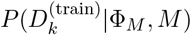 is the probability that 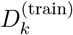 is generated by simulating model *M* with Φ_*M*_ (see Eq. 12), and 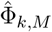 is the set of estimated parameters that maximizes that probability. For optimization, we used a combination of gradient-free (Subplex^109^; for a broad search of the parameter space) and gradient-based optimization algorithms (L-BFGS^110^; for fine-tuning), starting from 5 differently chosen initial conditions (see Code and data availability).

We then evaluated all models on the testing set: For each participant *n* in the testing set 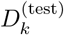 of fold *k*, we evaluated the cross-validated log-likelihood as

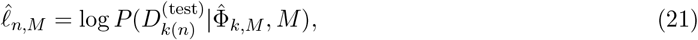

where 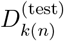 is the data of participant *n* (which we assumed to be in the testing set of fold *k*). We then used the cross-validated log-likelihoods in the Bayesian model selection method of ref. ^67^ with the random effects assumption: We assumed that, with an unknown probability *P*_*M*_, the data of each participant *n* was generated by simulating model *M*_*n*_ = *M*. The goal of the model comparison is to infer probability *P*_*M*_ for all models; the one with the highest *P*_*M*_ is the most probable model of most participants. To do so, we performed Markov Chain Monte Carlo sampling (using Metropolis Hasting algorithm ^54^ with 100 chains of length 10^′^000) and estimated the joint posterior distribution over *P*_novelty_, *P*_inf-gain_, *P*_surprise_, and *P*_opt. init._. Fig. 4B shows the expected value of *P*_*M*_ (the expected posterior probability; Fig. 4B1) and the probability of *P*_*M*_ being higher than *P*_*M*_^′^ for all *M* ^′^ ≠ *M* (the protected exceedance probabilities; Fig. 4B2). Fig. 4E shows the protected exceedance probabilities when the posterior distribution is evaluated conditioned on participants’ data in only one of the reward groups. See ref. ^37,79^for a similar approach and ref. ^41,67,68^for tutorials on the topic.

Finally, for each participant *n* in the testing set 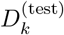 of fold *k*, we evaluated the accuracy rate of novelty-seeking (Fig. 4) in predicting the participant’s actions (conditioned on the past actions) in each episode, i.e., we evaluated the ratio of actions where novelty-seeking with parameter 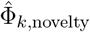 assigned the highest probability to the participant’s chosen action; whenever the maximum probability was shared between 2 or 3 actions, we considered the prediction 1/2 or 1/3 correct, respectively (i.e., a random model would have a 33% accuracy rate).

### Posterior predictive checks and model-recovery

For each model *M* ∈ {novelty, inf-gain, surprise, opt. init.}, we repeated the following steps 1500 times: 1. We picked, with one-third probability, the fitted parameter 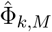 of fold *k* ∈ {1, 2, 3}. 2. We picked, with one-third probability, one of the reward conditions (i.e., 2 CHF, 3 CHF, and 4 CHF). 3. We simulated model *M* with parameters 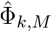 for 5 episodes in our environment, i.e., we sampled a trajectory *D* from 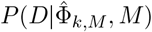 (with the *G*^*^ of the environment corresponding to the reward group picked in step 2). As a result, we ended up with 1500 simulated agents (with randomly sampled parameters) for each algorithm.

Depending on their exploration strategy and parameters, some simulated agents kept exploring the stochastic part of the environment and did not escape it. Hence, we stopped simulations of each episode after 3000 actions; note that the median number of actions taken by human participants is less than 100 (Fig. 2B-C). Accordingly, we considered the simulated agents who took more than 3000 actions in any of the 5 episodes to be similar to the human participants who quit the experiment and excluded them from further analyses. Moreover, we applied the same criterion that we used for the human participants and excluded, separately for each algorithm, the simulated agents who took more than 3 times the group-average number of actions in episodes 2-5 to finish the experiment. We then analyzed the remaining simulated agents. Fig. 2D-F shows the data statistics of simulated novelty-seeking agents compared to human participants.

Fig. 4C shows the median relative error (absolute difference divided by SE) of different algorithms in reproducing 44 group-level statistics: (1) Ratio of excluded agents, (2) number of actions in episode 1, (3-6) fractions of trials spent in trap states and stochastic parts during the 1st and 2nd halves of episode 1 (Fig. 2A), (7-10) median number actions in episodes 2-5 for each reward group and its correlation with reward value (Fig. 2B1), (11-14) fraction of trials spent in the stochastic part in episodes 2-5 for each reward group and its correlation with reward value (Fig. 2B2), (15-17) correlation of episode length with episode number for each reward group (e.g., Fig. 2C for the 2 CHF group), (18-20) correlation of the fraction of trials spent in the stochastic part with the episode number for each reward group, and (21-44) the ratio of taking different actions (2 possibilities, i.e., progressing action and self-looping/stochastic action) in different progressing states (3 possibilities, i.e., states 1-3, state 4, and states 5-6) and in different periods of the experiment (4 possibilities, i.e., episode 1 for all participants and episodes 2-5 separately for each reward group). See Supplementary Materials for details.

Finally, for the simulated data of each algorithm, we repeated our model selection procedure (i.e., 3-fold cross-validation plus Bayesian model selection) on the action choices of 5 groups of 60 simulated agents (20 from each participant group, i.e., 2 CHF, 3 CHF, and 4 CHF). We always successfully recovered the model that had generated the data, using almost the same number of simulated agents (60) as human participants (57). See insets in Fig. 4B for confusion matrices.

## Supporting information

Supplementary materials to Even If Suboptimal, Novelty Drives Human Exploration

## Acknowledgement

AM thanks Vasiliki Liakoni, Johanni Brea, Sophia Becker, Martin Barry, Valentin Schmutz, and Guil-laume Bellec for many useful discussions. All authors thank Peter Dayan and Joshua Gold for valuable feedback on the manuscript. This research was supported by the Swiss National Science Foundation No. CRSII2 147636 (Sinergia; MHH and WG), No. 200020 184615 (WG), and No. 200020 207426 (WG) and by the European Union Horizon 2020 Framework Program under grant agreement No. 785907 (Human Brain Project, SGA2; MHH and WG).

## Author Contributions

AM, HAX, MHH, and WG developed the study concept and designed the experiment. HAX and WL conducted the experiment and collected the data. AM designed the algorithms, did the formal analyses, and analyzed the data. AM, MHH, and WG wrote the paper.

## Code and data availability

All code and data needed to reproduce the results reported in this manuscript will be made publicly available after publication acceptance.

