## Supplementary materials to Even If Suboptimal, Novelty Drives Human Exploration for "Even if suboptimal, novelty drives human exploration"

### 1 Supplementary Results

#### 1.1 Different intrinsic rewards show different attraction to the stochastic part

To confirm that the stochastic part of our experimental paradigm preserves the essential features of a noisy TV (Aubret et al., 2023; Burda et al., 2019), we simulated three groups of RL agents (500 agents per group) exploring our environment by seeking (i) surprise, (ii) novelty, or (iii) information gain. As a control group, we also simulated 500 random agents taking each action with an equal probability of 1/3.

We aimed to quantify the isolated effect of the intrinsic reward on the simulated agents' attraction to the stochastic part. To achieve this, we considered the most efficient version of each exploration strategy by employing the model-based branch of our algorithm in Algorithm 1. Additionally, we removed the extrinsic-reward-seeking component of the algorithm; this would allow us to analyze the purely exploratory behavior of the RL agents in isolation (similar to Burda et al. (2019); Pathak et al. (2017)).

The resulting algorithm included a total of 7 parameters: $\{\lambda_{\text{int}}, \kappa, \epsilon_{\text{new}}, \epsilon_{\text{obs}}, T_{PS,\text{int}}, \beta^{(1)}_{\text{MB,int}}, \beta^{(2)}_{\text{MB,int}}\}$. To avoid arbitrariness in parameter selection, we assumed perfect model-building by setting $\kappa = 1$ and nearly perfect planning by setting $T_{PS,\text{int}} = 100$. Furthermore, we chose the discount factor $\lambda_{\text{int}}$ and the prior parameters $\epsilon_{\text{new}}$ and $\epsilon_{\text{obs}}$ based on the range of fitted parameters reported by Xu et al. (2021): $\lambda_{\text{int}} = 0.70$, $\epsilon_{\text{new}} = 10^{-5}$, and $\epsilon_{\text{obs}} = 10^{-4}$. Finally, after fixing these parameter values, we fine-tuned $\beta^{(1)}_{\text{MB,int}}$ to minimize the average length of episode 1; in other words, we fined-tuned $\beta^{(1)}_{\text{MB,int}}$ such that the agents would find the goal as fast as possible. We then simulated the agents for an additional 5 episodes with $\beta^{(2)}_{\text{MB,int}} = \beta^{(1)}_{\text{MB,int}}$. Each episode ended when agents reached the goal state $G^*$, even though no extrinsic reward was associated with $G^*$. Depending on their exploration strategy, some simulated agents kept exploring the stochastic part of the environment and did not escape it. Hence, we stopped simulations of each episode after 3000 actions.

We characterized the exploratory behavior of different agents during episodes 2–5 by measuring the search duration (Fig. S1A) and the fraction of trials spent in the stochastic part (Fig. S1B). For agents seeking information gain, both the search duration and the fraction of trials in the stochastic part decreased over episodes (Fig. S1A3 and B3). Conversely, novelty- and surprise-seeking agents exhibited the opposite pattern (Fig. S1A1-A2 and B1-B2). Notably, surprise-seeking agents were often (i.e., in $> 50\%$ of simulations in episode 5) stuck in the stochastic part, failing to escape within 3000 actions (Fig. S1A1). By design, random agents exhibited consistent behavior across episodes (Fig. S1A4 and B4): They had a persistently higher search duration, compared to novelty- and information-gain-seeking agents (Fig. S1A4 versus Fig. S1A2-A3), but spent only a marginal fraction of their time in the stochastic part of the environment (Fig. S1B4 versus Fig. S1B1-B3).

These findings confirm that the stochastic part of our experimental paradigm effectively replicates the distinct exploration patterns previously associated with different intrinsic rewards (Aubret et al., 2023).

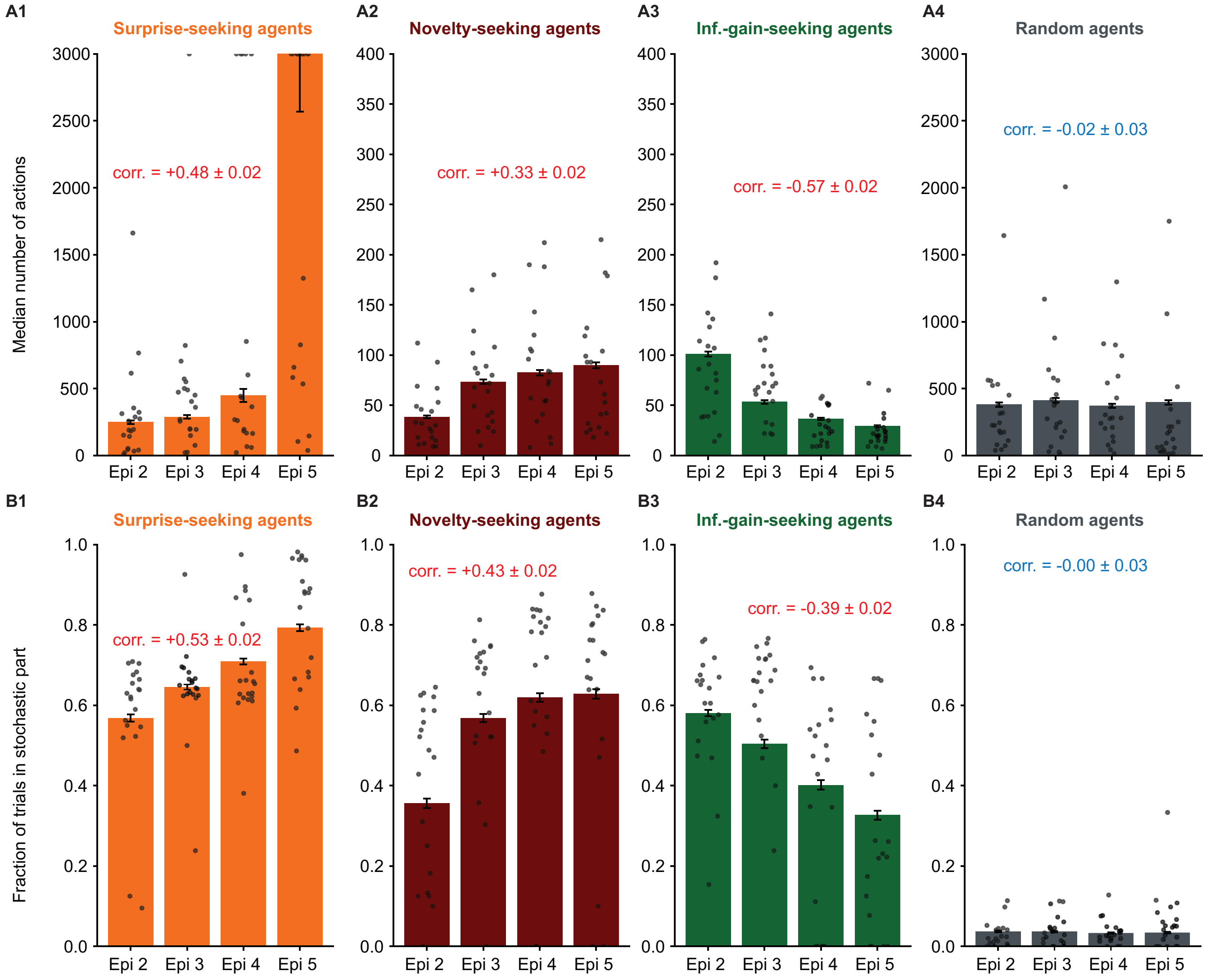

Figure S1: **Efficient exploration driven by different intrinsic rewards shows different patterns of attraction to the stochastic part (Supplementary to Figure 1 in the main text). A.** Median number of actions in episodes 2-5 for agents simulated by different algorithms (500 simulations for each algorithm). Error bars show the SEMed. Correlations denote the average (across simulations) Pearson correlation between the number of actions and the episode number. **B.** Fraction of trials spent in the stochastic part in episodes 2-5. Error bars show the SEMean. In all panels, single dots show the data of (20 out of 500) individual simulations. Correlations denote the average (across simulations) Pearson correlation between the fraction of trials in the stochastic part and the episode number.

#### 1.2 Detailed summary statistics for posterior predictive checks

For model selection based on Posterior Predictive Checks (PPC), we reported the median relative error of different algorithms in reproducing 44 summary statistics of human data (Figure 4C in the main text). Fig. S2 displays the relative error for all summary statistics separately. Consistent with our conclusion in the main text, novelty-seeking most accurately reproduces the majority of the summary statistics of human action choices (Fig. S2; 1st row versus the others).

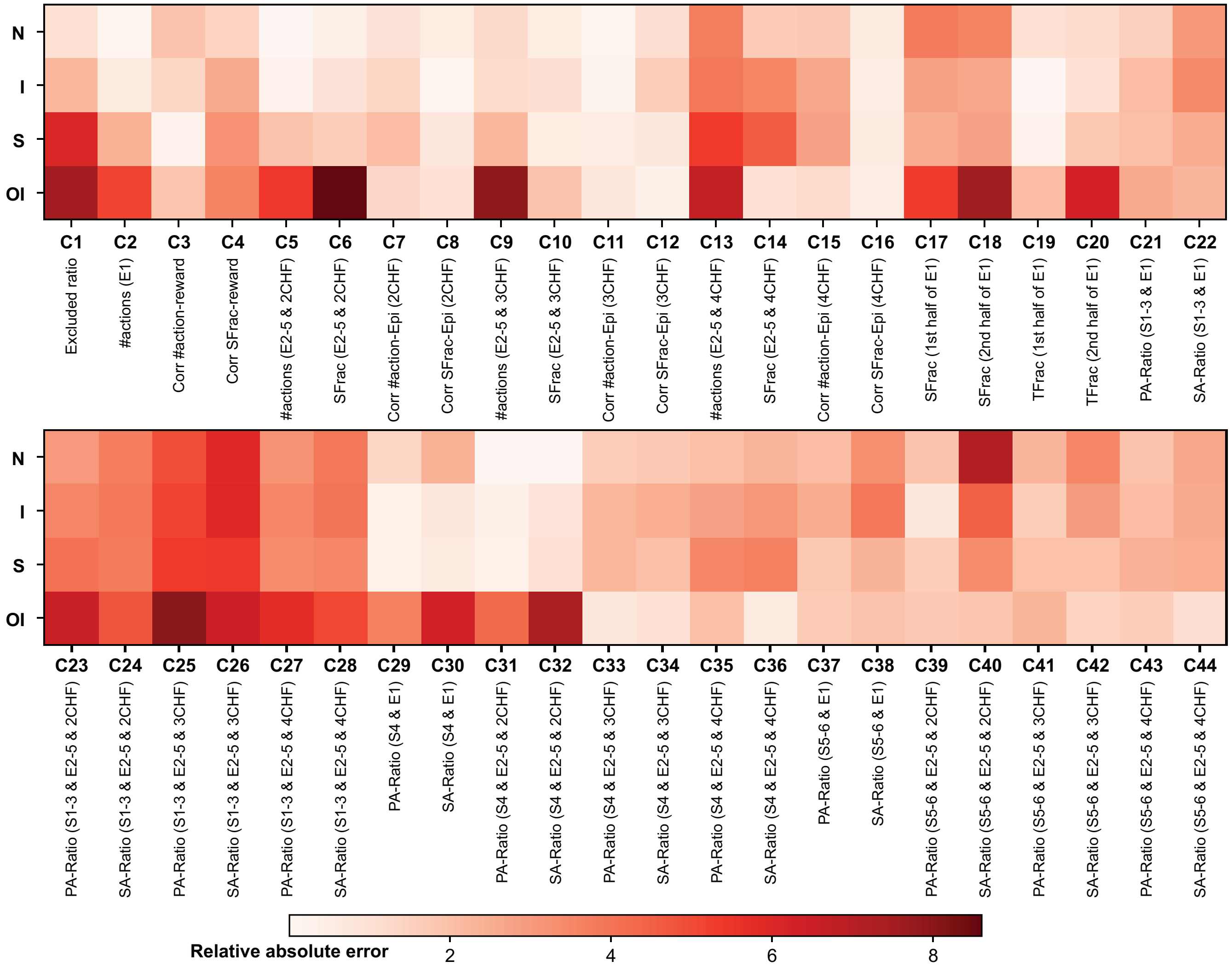

Figure S2: **Relative error of different algorithms in replicating 44 summary statistics of human data (Supplementary to Figure 4C in the main text).** The heatmap shows the relative error evaluated as the absolute difference in the summary statistics, divided by their standard errors. Each row corresponds to one of the four algorithms: novelty-seeking (N), information-gain-seeking (I), surprise-seeking (S), and exploration based on optimistic initialization (OI). Each column (C1-C44) corresponds to one of the 44 summary statistics: the ratio of excluded agents (C1); the median number of actions in episode 1 (C2); correlation between the reward value of $G^*$ and the number of actions in episodes 2-5 (C3) or the fraction of trials spent in the stochastic states in episodes 2-5 (C4); the median number of actions (C5, C9, and C13) and the average fraction of time in the stochastic states (C6, C10, and C14) during episodes 2-5 for different reward groups (2CHF, 3CHF, and 4CHF), respectively; correlation between the episode number and the number of actions (C7, C11, and C15) or the fraction of time in the stochastic states (C8, C12, and C16) during episodes 2-5 for different reward groups (2CHF, 3CHF, and 4CHF), respectively; fraction of trials spent in the stochastic part (C17 and C18) and the trap states (C19 and C20) during the two parts (the 1st and 2nd halves) of episode 1, respectively; the ratio (C21-C44) of taking different actions (2 possibilities, i.e., progressing action and self-looping/stochastic action, abbreviated by PA and SA, respectively) in different progressing states (3 possibilities, i.e., states 1-3, state 4, and states 5-6, abbreviated by S1-3, S4, and S5-6) and in different periods of the experiment (4 possibilities, i.e., episode 1 for all participants, abbreviated by E1, and episodes 2-5 separately for each reward group, indexed by E2-5 & 2CHF, 3CHF, or 4CHF, respectively).

### 2 Supplementary Methods

#### 2.1 Model-building in an environment of unknown size

We use ideas from non-parametric Bayesian inference (Gershman and Blei, 2012; Ghahramani, 2013) and Dirichlet processes (Teh, 2010) to derive a Bayesian estimate $p^{(t)}(s'|s,a)$ of the transition probabilities in an environment of unknown size.

##### 2.1.1 Time dependent base distribution as the expected prior

Consider the 1st time an agent takes action $a$ at state $s$. Which is the state $s'$ where the agent is expected to visit next given that it has zero experience for taking action $a$ at state $s$? There are two possibilities: (i) $s'$ is one of the already known state, i.e., $s' \in \mathcal{S}^{(t)}$, and (ii) $s'$ is one of the infinitely many imaginable states $\mathcal{S}$ that the agent has not observed yet, i.e., $s' \notin \mathcal{S}^{(t)}$. We assume that the agent considers different weights for these two possibilities even in the prior distribution. We give a precise definition of this prior distribution in the next subsection, but we first need to define our *time dependent base distribution* (Teh, 2010) which we will use later.

We define the probability measure $H$ as a continuous probability distribution (i.e. without any atom) on the space of all imaginable states $\mathcal{S}$ – e.g., the space of all images that can appear on the computer screen. Our results are independent of the exact shape of $H$ – as long as it is a *continuous* probability distribution. We then define the time-dependent base distribution on $\mathcal{S}$ as

$$H^{(t)} := \frac{\epsilon_{\text{new}}}{\epsilon_{\text{new}} + \epsilon_{\text{obs}}|\mathcal{S}^{(t)}|} H + \frac{\epsilon_{\text{obs}}}{\epsilon_{\text{new}} + \epsilon_{\text{obs}}|\mathcal{S}^{(t)}|} \sum_{s \in \mathcal{S}^{(t)}} \delta_s, \tag{S1}$$

where $\delta_s$ is the Dirac measure at $s$, $\epsilon_{\text{obs}}$ and $\epsilon_{\text{new}}$ are the weights for combining the two possibilities of (i) transiting to a known state $s' \in \mathcal{S}^{(t)}$ and (ii) transiting to a new and unknown state $s' \in \mathcal{S}$. In the next section, we use this base distribution as $p^{(t)}(.|s,a)$ for any state-action pair $(s,a)$ that has not been experienced before.

##### 2.1.2 Derivation of the world-model

We indicate the matrix of transition probabilities as a parameter $\Theta$ that fully summarizes the environment. Then, given underlying $\Theta = \theta : \mathcal{S} \times \mathcal{A} \to \text{Measures}[\mathcal{S}]$, we have

$$\mathbb{P}(S_{t+1} = s'|S_t = s, A_t = a, \Theta = \theta) := \theta_{s,a}(s') \tag{S2}$$

for any $s$ and $s' \in \mathcal{S}$ and $a \in \mathcal{A}$. Given the sequence of states $S_{1:t} = s_{1:t}$ and actions $A_{1:t-1} = a_{1:t-1}$, an agent's belief about the transition matrix $\theta$ is defined as the posterior

$$q^{(t)}(\theta) := \mathbb{P}^{(t)}(\theta|s_{1:t}, a_{1:t-1}) \propto \mathbb{P}^{(t)}(\theta)\mathbb{P}^{(t)}(s_{2:t}|\theta, a_{1:t-1}, s_1) = \mathbb{P}^{(t)}(\theta) \prod_{t'=1}^{t-1} \theta_{s_{t'},a_{t'}}(s_{t'+1}), \tag{S3}$$

where the prior $\mathbb{P}^{(t)}$ is a time-dependent prior distribution over transition probabilities. We assume that $\Theta_{s,a}$s are *a priory* i.i.d. samples of a Dirichlet process prior (Teh, 2010) with the base distribution $H^{(t)}$ and a time-dependent concentration parameter $\alpha^{(t)}$, that is, for any finite and countable $\mathcal{S}' \subseteq \mathcal{S}$,

$$\mathbb{P}^{(t)}\big(\{\theta_{s,a} : s \in \mathcal{S}', a \in \mathcal{A}\}\big) = \prod_{s \in \mathcal{S}', a \in \mathcal{A}} \text{DP}(\theta_{s,a}; \alpha^{(t)}, H^{(t)}), \tag{S4}$$

where DP stands for Dirichlet Process. $H^{(t)}$ is the prior expected value of $\Theta$ and can be seen as a prior estimate of transition probabilities, and $\alpha^{(t)}$ shows how many samples this estimate is worth (Efron and

Hastie, 2016; Teh, 2010). Putting a weight of $\epsilon_{\text{obs}}$ for each known state and $\epsilon_{\text{new}}$ for all unknown state (Eq. S1), we end up with $\alpha^{(t)} = \epsilon_{\text{new}} + \epsilon_{\text{obs}}|\mathcal{S}^{(t)}|$ as the number of samples that $H^{(t)}$ is worth.

It is straightforward to show that the posterior distribution $q^{(t)}$ has the same form as the prior (Teh, 2010), that is, for any finite and countable $\mathcal{S}' \subseteq \mathcal{S}$,

$$q^{(t)}\big(\{\theta_{s,a} : s \in \mathcal{S}', a \in \mathcal{A}\}\big) = \prod_{s\in\mathcal{S}',a\in\mathcal{A}} \text{DP}\Big(\theta_{s,a}\,;\, \alpha^{(t)} + C^{(t)}_{s,a}\,,\, \frac{\alpha^{(t)}}{\alpha^{(t)} + C^{(t)}_{s,a}} H^{(t)} + \frac{1}{\alpha^{(t)} + C^{(t)}_{s,a}} \sum_{s'\in\mathcal{S}^{(t)}} C^{(t)}_{s,a,s'}\delta_{s'}\Big), \tag{S5}$$

where $C^{(t)}_{s,a}$ is the number of times action $a$ has been taken at state $s'$ until time $t$, and $C^{(t)}_{s,a,s'}$ is the number of times transition $(s,a) \to s'$ has been experienced. We consider the posterior expected value of $\Theta$ as an estimate of the world-model (Teh, 2010)

$$\begin{aligned} p^{(t)}(s'|s,a) := \hat{\theta}^{(t)}_{s,a}(s') := \mathbb{E}_{q^{(t)}}[\Theta_{s,a}(s')] &= \frac{\alpha^{(t)}}{\alpha^{(t)} + C^{(t)}_{s,a}} H^{(t)}(s') + \frac{1}{\alpha^{(t)} + C^{(t)}_{s,a}} \sum_{s''\in\mathcal{S}^{(t)}} C^{(t)}_{s,a,s''}\delta_{s''}(s') \\ &= \frac{\alpha^{(t)}(1-c_t)}{\alpha^{(t)} + C^{(t)}_{s,a}} H(s') + \frac{1}{\alpha^{(t)} + C^{(t)}_{s,a}} \sum_{s''\in\mathcal{S}^{(t)}} \Big(\frac{\alpha^{(t)}c_t}{|\mathcal{S}^{(t)}|} + C^{(t)}_{s,a,s''}\Big)\delta_{s''}(s'), \end{aligned} \tag{S6}$$

where we used $c_t = \frac{\epsilon_{\text{obs}}|\mathcal{S}^{(t)}|}{\epsilon_{\text{new}}+\epsilon_{\text{obs}}|\mathcal{S}^{(t)}|}$ to shorten the notation. Eq. S6 can be simplified and written as

$$p^{(t)}(s'|s,a) = \hat{\theta}^{(t)}_{s,a}(s') = \begin{cases} \frac{\epsilon_{\text{obs}}+C^{(t)}_{s,a,s'}}{\epsilon_{\text{new}}+\epsilon_{\text{obs}}|\mathcal{S}^{(t)}|+C^{(t)}_{s,a}} & \text{if} \quad s' \in \mathcal{S}^{(t)}\,, \\ \frac{\epsilon_{\text{new}}}{\epsilon_{\text{new}}+\epsilon_{\text{obs}}|\mathcal{S}^{(t)}|+C^{(t)}_{s,a}} & \text{if} \quad s' = s_{\text{new}}\,. \end{cases} \tag{S7}$$

where by $s' = s_{\text{new}}$ we mean $s' \notin \mathcal{S}^{(t)}$, i.e.,

$$\hat{\theta}^{(t)}_{s,a}(s_{\text{new}}) := \mathbb{E}_{\Theta_{s,a}\sim q^{(t)}}\Big[\mathbb{E}_{S'\sim\Theta_{s,a}}[\mathcal{I}_{S'\notin\mathcal{S}^{(t)}}]\Big] = \frac{\alpha^{(t)}(1-c_t)}{\alpha^{(t)} + C^{(t)}_{s,a}} \int_{s'\notin\mathcal{S}^{(t)}} H(s')ds' = \frac{\alpha^{(t)}(1-c_t)}{\alpha^{(t)} + C^{(t)}_{s,a}}. \tag{S8}$$

For the case of $\epsilon_{\text{new}} = 0$, $\epsilon_{\text{obs}} = \epsilon$, and $\mathcal{S}^{(t)} = \mathcal{S}$ being a finite and countable set, the transition matrix is the same as the transition matrix conventionally used for finite state-spaces (Kolter and Ng, 2009; Xu et al., 2021). For the case of $\epsilon_{\text{obs}} = 0$, the transition matrix has the form of a Chinese restaurant process (Blei and Frazier, 2011; Teh, 2010).

To account for imperfect model-building, we use leaky counts $\tilde{C}^{(t)}_{s,a,s'}$ and $\tilde{C}^{(t)}_{s,a} = \sum_{s'} \tilde{C}^{(t)}_{s,a,s'}$ instead of $C^{(t)}_{s,a}$ and $C^{(t)}_{s,a,s'}$, where $\tilde{C}^{(t)}_{s,a,s'}$ is recursively updated via

$$\tilde{C}^{(t+1)}_{s,a,s'} = \begin{cases} \kappa\tilde{C}^{(t)}_{s,a,s'} + \delta_{s',s_{t+1}} & \text{if}\, s = s_t\,,\; a = a_t \\ \tilde{C}^{(t)}_{s,a,s'} & \text{otherwise,} \end{cases} \tag{S9}$$

where $\delta$ is the Kronecker delta function, $\tilde{C}^{(0)}_{s,a,s'} = 0$, and $\kappa \in [0,1]$ is the leak parameter; this is a common modeling choice in neuroscience and psychology (Liakoni et al., 2021; Meyniel et al., 2016; Yu and Cohen, 2009). If $\kappa = 1$, then $\tilde{C}^{(t+1)}_{s,a,s'} = C^{(t+1)}_{s,a,s'}$.

One may argue that the whole Bayesian formulation could be avoided by considering Eq. S7 as the starting point – similar to how we present the model in Methods in the main text. However, as we will see in the next two sections, Eq. S7 without the Bayesian formulation of this section is not enough for deriving (i) the update rule for the model-based branch and (ii) evaluating information gain.

#### 2.2 Prioritized sweeping for updating the MB Q-values

Given a reward function $R$ (i.e., $R_{\text{ext}}$ for the extrinsic reward or $R_{\text{int},t}$ for the intrinsic reward) the Bellman equations (Puterman, 1994; Sutton and Barto, 2018) are

$$Q^{(t)}(s,a) = \mathbb{E}_{S' \sim \hat{\theta}^{(t)}_{s,a}} \Big[ R_{s,a}(S') + \lambda \max_{a' \in \mathcal{A}} Q^{(t)}(S', a') \Big], \tag{S10}$$

where $Q^{(t)}$ is the Q-value (i.e., $Q^{(t)}_{\text{MB,ext}}$ for the extrinsic reward or $Q^{(t)}_{\text{MB,int}}$ for the intrinsic reward), $\lambda \in [0,1)$ is the discount factor (i.e., $\lambda_{\text{ext}}$ for the extrinsic reward or $\lambda_{\text{int}}$ for the intrinsic reward), and we show $R(s, a \to s')$ by $R_{s,a}(s')$ to shorten the notation.

We assume that $R_{s,a}(s')$ is the same for all $s' \notin \mathcal{S}^{(t)}$. Then, given the fact that $\hat{\theta}^{(t)}_{s,a}(s')$ is also the same for all $s' \notin \mathcal{S}^{(t)}$ or $s \notin \mathcal{S}^{(t)}$ (Eq. S7), $Q^{(t)}(s,a)$ takes the same value for all $s \notin \mathcal{S}^{(t)}$ and for all actions $a \in \mathcal{A}$. Hence, Eq. S10 can be re-written as

$$Q^{(t)}(s,a) = \sum_{s' \in \mathcal{S}^{(t)}} \hat{\theta}^{(t)}_{s,a}(s') \Big( R_{s,a}(s') + \lambda V^{(t)}(s') \Big) + \hat{\theta}^{(t)}_{s,a}(s_{\text{new}}) \Big( R_{s,a}(s_{\text{new}}) + \lambda V^{(t)}(s_{\text{new}}) \Big), \tag{S11}$$

where $V^{(t)}(s') := \max_{a' \in \mathcal{A}} Q^{(t)}(s', a')$. We use the observation that $V^{(t)}(s_{\text{new}}) = Q^{(t)}(s_{\text{new}}, a)$ is independent of $a$ and find $V^{(t)}(s_{\text{new}})$ by solving

$$\begin{aligned} V^{(t)}(s_{\text{new}}) &= \sum_{s' \in \mathcal{S}^{(t)}} \hat{\theta}^{(t)}_{s_{\text{new}}}(s') \Big( R_{s_{\text{new}}}(s') + \lambda V^{(t)}(s') \Big) + \hat{\theta}^{(t)}_{s_{\text{new}}}(s_{\text{new}}) \Big( R_{s_{\text{new}}}(s_{\text{new}}) + \lambda V^{(t)}(s_{\text{new}}) \Big) \\ &= \frac{\epsilon_{\text{obs}}}{\epsilon_{\text{new}} + \epsilon_{\text{obs}} |\mathcal{S}^{(t)}|} \sum_{s' \in \mathcal{S}^{(t)}} \Big( R_{s_{\text{new}}}(s') + \lambda V^{(t)}(s') \Big) + \frac{\epsilon_{\text{new}}}{\epsilon_{\text{new}} + \epsilon_{\text{obs}} |\mathcal{S}^{(t)}|} \Big( R_{s_{\text{new}}}(s_{\text{new}}) + \lambda V^{(t)}(s_{\text{new}}) \Big), \end{aligned} \tag{S12}$$

where we also used the observation that $R_{s_{\text{new}}}(s') := R_{s_{\text{new}},a}(s')$ is independent of $a$. The solution to Eq. S12 is given by

$$\begin{aligned} V^{(t)}(s_{\text{new}}) &= \frac{\epsilon_{\text{obs}}}{(1-\lambda)\epsilon_{\text{new}} + \epsilon_{\text{obs}} |\mathcal{S}^{(t)}|} \sum_{s' \in \mathcal{S}^{(t)}} \Big( R_{s_{\text{new}}}(s') + \lambda V^{(t)}(s') \Big) + \frac{\epsilon_{\text{new}}}{(1-\lambda)\epsilon_{\text{new}} + \epsilon_{\text{obs}} |\mathcal{S}^{(t)}|} R_{s_{\text{new}}}(s_{\text{new}}) \\ &= W^{(t)}_{\text{obs}} \sum_{s' \in \mathcal{S}^{(t)}} \Big( R_{s_{\text{new}}}(s') + \lambda V^{(t)}(s') \Big) + W^{(t)}_{\text{new}} R_{s_{\text{new}}}(s_{\text{new}}), \end{aligned} \tag{S13}$$

where in the last line we shortened the notation by defining constants

$$W^{(t)}_{\text{obs}} := \frac{\epsilon_{\text{obs}}}{(1-\lambda)\epsilon_{\text{new}} + \epsilon_{\text{obs}} |\mathcal{S}^{(t)}|} \quad \text{and} \quad W^{(t)}_{\text{new}} := \frac{\epsilon_{\text{new}}}{(1-\lambda)\epsilon_{\text{new}} + \epsilon_{\text{obs}} |\mathcal{S}^{(t)}|}. \tag{S14}$$

We can combine Eq. S11 and Eq. S13 to derive a set of equations for the $Q$-values of states in $\mathcal{S}^{(t)}$:

$$\begin{aligned} Q^{(t)}(s,a) =& \lambda \sum_{s' \in \mathcal{S}^{(t)}} \Big( \hat{\theta}^{(t)}_{s,a}(s') + \lambda \hat{\theta}^{(t)}_{s,a}(s_{\text{new}}) W^{(t)}_{\text{obs}} \Big) V^{(t)}(s') + \sum_{s' \in \mathcal{S}^{(t)}} \hat{\theta}^{(t)}_{s,a}(s') R_{s,a}(s') + \\ & \hat{\theta}^{(t)}_{s,a}(s_{\text{new}}) \Big( R_{s,a}(s_{\text{new}}) + \lambda W^{(t)}_{\text{new}} R_{s_{\text{new}}}(s_{\text{new}}) + \lambda W^{(t)}_{\text{obs}} \sum_{s' \in \mathcal{S}^{(t)}} R_{s_{\text{new}}}(s') \Big). \end{aligned} \tag{S15}$$

To solve this set of equations, we use prioritized sweeping (Brea, 2017; Sutton and Barto, 2018; Van Seijen and Sutton, 2013) with some modifications (similar to Xu et al. (2021)). The modified algorithm is presented in Alg. 4 in which we use the $Q$Update operator defined as

$$
\begin{aligned}
&Q\text{Update}(s, a; \lambda, \hat{\theta}, V, R, W_{\text{obs}}, W_{\text{new}}, \tilde{\mathcal{S}}) \\
&\qquad := \lambda \sum_{s' \in \tilde{\mathcal{S}}} \Big(\hat{\theta}_{s,a}(s') + \lambda \hat{\theta}_{s,a}(s_{\text{new}}) W_{\text{obs}}\Big) V(s') + \sum_{s' \in \tilde{\mathcal{S}}} \hat{\theta}_{s,a}(s') R_{s,a}(s') + \\
&\qquad\quad \hat{\theta}_{s,a}(s_{\text{new}}) \Big(R_{s,a}(s_{\text{new}}) + \lambda W_{\text{new}} R_{s_{\text{new}}}(s_{\text{new}}) + \lambda W_{\text{obs}} \sum_{s' \in \tilde{\mathcal{S}}} R_{s_{\text{new}}}(s')\Big).
\end{aligned}
\tag{S16}
$$

#### 2.3 Derivation of information gain

Information-gain-seeking algorithms (Mobin et al., 2014; Poli et al., 2024; Schmidhuber, 2010) consider the intrinsic reward as the amount of change in the world-model $\hat{\theta}^{(t)}_{s,a}$ upon observing the transition $(s, a) \to s'$, often defined as

$$
R_{\text{int},t}(s, a \to s') = IG^{(t)}(s, a \to s') = \mathrm{D}_{\text{KL}}\Big[\ \hat{\theta}^{(t)}_{s,a} \ ||\ \hat{\theta}^{(t+1)}_{s,a\to s'}\ \Big], \tag{S17}
$$

where $\hat{\theta}^{(t+1)}_{s,a\to s'}$ is $\hat{\theta}^{(t+1)}_{s,a}$ if $S_{t+1} = s'$, and $\mathrm{D}_{\text{KL}}$ is the Kullback-Leibler divergence (Cover, 1999). In different contexts, $IG^{(t)}(s, a \to s')$ is also called Postdictive surprise (Kolossa et al., 2015), but it has a fundamentally different behavior from the *prediction* surprise $-\log \hat{\theta}^{(t)}_{s,a}(s')$ that we used for our surprise-seeking algorithm (see Methods in the main text and Modirshanechi et al. (2022) for more discussion).

If $s' \notin \mathcal{S}_t$, the naïve definition of $\mathrm{D}_{\text{KL}}$ cannot be used in Eq. S17 because $\hat{\theta}^{(t)}_{s,a}$ and $\hat{\theta}^{(t+1)}_{s,a\to s'}$ has different supports for their atoms. To resolve this issue, Mobin et al. (2014) propose a padding mechanism as a heuristic solution. We use a more general definitions of $\mathrm{D}_{\text{KL}}$ as the expected Radon–Nikodym derivative of $\hat{\theta}^{(t)}_{s,a}$ with respect to $\hat{\theta}^{(t+1)}_{s,a\to s'}$ that is well-defined in our Bayesian framework:

$$
\mathrm{D}_{\text{KL}}\Big[\ \hat{\theta}^{(t)}_{s,a} \ ||\ \hat{\theta}^{(t+1)}_{s,a\to s'}\ \Big] = \mathbb{E}_{S'' \sim \hat{\theta}^{(t)}_{s,a}}\left[\frac{d\hat{\theta}^{(t)}_{s,a}}{d\hat{\theta}^{(t+1)}_{s,a\to s'}}(S'')\right], \tag{S18}
$$

where $\frac{d\hat{\theta}^{(t)}_{s,a}}{d\hat{\theta}^{(t+1)}_{s,a\to s'}}(S'')$ is the Radon–Nikodym derivative of $\hat{\theta}^{(t)}_{s,a}$ with respect to $\hat{\theta}^{(t+1)}_{s,a\to s'}$ at $S''$; we note that $\hat{\theta}^{(t)}_{s,a}$ is always absolutely continuous with respect to $\hat{\theta}^{(t+1)}_{s,a\to s'}$, which implies that the Radon–Nikodym derivative is well-defined in our case. Accordingly, for $s' \in \mathcal{S}^{(t)}$, the Radon–Nikodym derivative is

$$
\frac{d\hat{\theta}^{(t)}_{s,a}}{d\hat{\theta}^{(t+1)}_{s,a\to s'}}(s'') = \begin{cases} \frac{\epsilon_{\text{new}} + \epsilon_{\text{obs}}|\mathcal{S}^{(t)}| + C^{(t)}_{s,a} + 1}{\epsilon_{\text{new}} + \epsilon_{\text{obs}}|\mathcal{S}^{(t)}| + C^{(t)}_{s,a}} & \text{if} \quad s'' \neq s', \\ \frac{\epsilon_{\text{new}} + \epsilon_{\text{obs}}|\mathcal{S}^{(t)}| + C^{(t)}_{s,a} + 1}{\epsilon_{\text{new}} + \epsilon_{\text{obs}}|\mathcal{S}^{(t)}| + C^{(t)}_{s,a}} \frac{\epsilon_{\text{obs}} + C^{(t)}_{s,a,s'}}{\epsilon_{\text{obs}} + C^{(t)}_{s,a,s'} + 1} & \text{if} \quad s'' = s'. \end{cases} \tag{S19}
$$

and for $s' \notin \mathcal{S}^{(t)}$, the Radon–Nikodym derivative is

$$
\frac{d\hat{\theta}^{(t)}_{s,a}}{d\hat{\theta}^{(t+1)}_{s,a\to s'}}(s'') = \begin{cases} \frac{\epsilon_{\text{new}} + \epsilon_{\text{obs}}|\mathcal{S}^{(t)}| + \epsilon_{\text{obs}} + C^{(t)}_{s,a} + 1}{\epsilon_{\text{new}} + \epsilon_{\text{obs}}|\mathcal{S}^{(t)}| + C^{(t)}_{s,a}} & \text{if} \quad s'' \neq s', \\ 0 & \text{if} \quad s'' = s'. \end{cases} \tag{S20}
$$

As a result, the information gain in Eq. S17 can be calculated as

$$R_{\mathrm{int},t}(s,a\to s') = \begin{cases} \log \frac{\epsilon_{\mathrm{new}}+\epsilon_{\mathrm{obs}}|\mathcal{S}^{(t)}|+C^{(t)}_{s,a}+1}{\epsilon_{\mathrm{new}}+\epsilon_{\mathrm{obs}}|\mathcal{S}^{(t)}|+C^{(t)}_{s,a}} + \hat{\theta}^{(t)}_{s,a}(s') \log \frac{\epsilon_{\mathrm{obs}}+C^{(t)}_{s,a,s'}}{\epsilon_{\mathrm{obs}}+C^{(t)}_{s,a,s'}+1} & \text{if} \quad s' \in \mathcal{S}^{(t)}\,, \\ \log \frac{\epsilon_{\mathrm{new}}+\epsilon_{\mathrm{obs}}|\mathcal{S}^{(t)}|+\epsilon_{\mathrm{obs}}+C^{(t)}_{s,a}+1}{\epsilon_{\mathrm{new}}+\epsilon_{\mathrm{obs}}|\mathcal{S}^{(t)}|+C^{(t)}_{s,a}} & \text{if} \quad s' \notin \mathcal{S}^{(t)}\,. \end{cases} \tag{S21}$$

If $\epsilon_{\mathrm{new}} \to 0$, then the momentary average gain in information after taking action $a$ in state $s$ can be written as

$$\begin{aligned} \bar{IG}^{(t)}(s,a) &:= \mathbb{E}_{S'\sim\hat{\theta}^{(t)}_{s,a}}\left[IG^{(t)}(s,a\to S')\right] \\ &= \log\left[1+\frac{1}{B^{(t)}_{s,a}}\right] - \sum_{s'\in\mathcal{S}^{(t)}} \left(\hat{\theta}^{(t)}_{s,a}(s')\right)^2 \log\left[1+\frac{1}{B^{(t)}_{s,a}\hat{\theta}^{(t)}_{s,a}(s')}\right], \end{aligned} \tag{S22}$$

where we defined $B^{(t)}_{s,a} := \epsilon_{\mathrm{obs}}|\mathcal{S}^{(t)}| + C^{(t)}_{s,a}$. With a few line of algebra, we can show

$$\frac{\partial \bar{IG}^{(t)}(s,a)}{\partial C^{(t)}_{s,a}} = -\frac{1}{B^{(t)}_{s,a}(1+B^{(t)}_{s,a})}\left[1 - \sum_{s'\in\mathcal{S}^{(t)}} \left(\hat{\theta}^{(t)}_{s,a}(s')\right)^2 \frac{1+B^{(t)}_{s,a}}{1+\hat{\theta}^{(t)}_{s,a}(s')B^{(t)}_{s,a}}\right] \leq 0, \tag{S23}$$

where the equality holds if an only if $\hat{\theta}^{(t)}_{s,a}(s') = 1$ for some $s'$. Hence, $\bar{IG}^{(t)}(s,a)$ is a decreasing function of the count $C^{(t)}_{s,a}$ of the state-action pair $(s,a)$, i.e., the more action $a$ is taken in state $s$, the less informative it becomes.

#### 2.4 Analysis of the MB optimistic initialization in episode 2

To theoretically analyze the influence of the MB optimistic initialization in episode 2, we make a few simplistic assumptions:

1. $\epsilon_{\mathrm{new}}$ in Eq. S7 is negligible.
2. All transition probabilities except for the ones between the stochastic states and the progressing action in state 6 (because of the only one-time experience) have been learned with certainty during the 1st episode.
3. The counts for the actions in the stochastic part are roughly the same for all states and actions, which we denote by $\bar{C}^{(t)}$, i.e., for any state $s_s$ in the stochastic part, we assume that $C^{(t)}_{s_s,a} = \bar{C}^{(t)}$ for every action $a$.

Given these assumptions, we have a symmetry between the stochastic states, implying that the $Q$-values in the stochastic part are the same for all states. Hence, in all the following equations, we use $s_s$ to denote a representative state in the stochastic part, use $a_p$ to refer to the progressing actions, use $a_s$ to the stochastic/self-looping actions, denote state 4 by $s4$, denote state 6 by $s6$, use $r_{G^*}$ to denote the reward value of the already discovered goal, and define

$$\bar{R} := \frac{1}{|\mathcal{S}^{(t)}|}(1+r_1^*+r_2^*) \quad \text{and} \quad \bar{V}^{(t)} := \frac{1}{|\mathcal{S}^{(t)}|}\sum_{s'} V^{(t)}_{\mathrm{MB,ext}}(s').$$

Using these notations and assumptions as well as Eq. S7 and Eq. S10, we have

$$\begin{aligned} Q^{(t)}_{\mathrm{MB,ext}}(s_s,a_p) &= p^{(t)}_s\left(\bar{R}+\lambda_{\mathrm{ext}}\bar{V}^{(t)}\right) + \lambda_{\mathrm{ext}}\left(1-p^{(t)}_s\right)V^{(t)}_{\mathrm{MB,ext}}(s4) \\ Q^{(t)}_{\mathrm{MB,ext}}(s_s,a_s) &= p^{(t)}_s\left(\bar{R}+\lambda_{\mathrm{ext}}\bar{V}^{(t)}\right) + \lambda_{\mathrm{ext}}\left(1-p^{(t)}_s\right)V^{(t)}_{\mathrm{MB,ext}}(s_s), \end{aligned} \tag{S24}$$

where

$$p_s^{(t)} = \frac{\epsilon_{\text{obs}}|\mathcal{S}^{(t)}|}{\epsilon_{\text{obs}}|\mathcal{S}^{(t)}| + \bar{C}^{(t)}}. \tag{S25}$$

Note that $\frac{p_s^{(t)}}{|\mathcal{S}^{(t)}|}$ is equal to the probability of transition to any state $s'$ for which $C_{s_s,a,s'} = 0$ (see Eq. S7).

If the optimal policy is to leave the stochastic part and go to the already discovered goal state, then we must have

$$\text{Condition 1: } Q^{(t)}_{\text{MB,ext}}(s_s, a_s) < Q^{(t)}_{\text{MB,ext}}(s_s, a_p) = V^{(t)}_{\text{MB,ext}}(s_s). \tag{S26}$$

According to Eq. S24, Condition 1 is equivalent to $Q^{(t)}_{\text{MB,ext}}(s_s, a_p) = V^{(t)}_{\text{MB,ext}}(s_s) \le V^{(t)}_{\text{MB,ext}}(s4)$, which, by using Eq. S24 again and after a few lines of algebra, can be written as

$$\text{Condition 1} \equiv \frac{p_s^{(t)}}{1 - \lambda_{\text{ext}}(1 - p_s^{(t)})}\Big[\bar{R} + \lambda_{\text{ext}}\bar{V}^{(t)}\Big] < V^{(t)}_{\text{MB,ext}}(s4). \tag{S27}$$

Given that the optimal policy under Condition 1 is to leave the stochastic part and go to the already discovered goal state, we can write the value of state 4 as

$$V^{(t)}_{\text{MB,ext}}(s4) = Q^{(t)}_{\text{MB,ext}}(s4, a_p) = \lambda_{\text{ext}}^2 Q^{(t)}_{\text{MB,ext}}(s6, a_p) = \lambda_{\text{ext}}^2\Big[\tilde{p}_g^{(t)}\big(\bar{R} + \lambda_{\text{ext}}\bar{V}^{(t)}\big) + \big(1 - p_g^{(t)}\big) r_{G^*}\Big], \tag{S28}$$

where

$$p_g^{(t)} = \frac{\epsilon_{\text{obs}}|\mathcal{S}^{(t)}|}{\epsilon_{\text{obs}}|\mathcal{S}^{(t)}| + 1} \quad \text{and} \quad \tilde{p}_g^{(t)} = p_g^{(t)} + \lambda_{\text{ext}}(1 - p_g^{(t)}). \tag{S29}$$

Note that $\frac{p_g^{(t)}}{|\mathcal{S}^{(t)}|}$ is equal to the probability of transition to any state $s'$ for which $C_{s6,a_p,s'} = 0$ (see Eq. S7). Using Eq. S28, we can simplify Eq. S27 as

$$\text{Condition 1} \equiv f_{\text{C1}}(r_1^*, r_2^*, \lambda_{\text{ext}}, \epsilon_{\text{obs}}, \bar{C}^{(t)}, |\mathcal{S}^{(t)}|) < r_{G^*}, \tag{S30}$$

with

$$f_{\text{C1}}(r_1^*, r_2^*, \lambda_{\text{ext}}, \epsilon_{\text{obs}}, \bar{C}^{(t)}, |\mathcal{S}^{(t)}|) := \frac{1}{\lambda_{\text{ext}}^2(1 - p_g^{(t)})}\Big[\frac{p_s^{(t)}}{1 - \lambda_{\text{ext}}(1 - p_s^{(t)})} - \lambda_{\text{ext}}^2 \tilde{p}_g^{(t)}\Big]\Big[\bar{R} + \lambda_{\text{ext}}\bar{V}^{(t)}\Big]. \tag{S31}$$

The variable $R^{(t)}_{\text{Stoch.}}$ in the Method section in the main text is $\lambda_{\text{ext}}^2 f_{\text{C1}}$.

An important observation is that, independently of the parameter values, we have

$$\lim_{\bar{C}^{(t)} \to \infty} f_{\text{C1}}(r_1^*, r_2^*, \lambda_{\text{ext}}, \epsilon_{\text{obs}}, \bar{C}^{(t)}, |\mathcal{S}^{(t)}|) < 0.$$

This implies that, for any value of $r_2^*$ and $r_{G^*} > 0$, increasing $\bar{C}^{(t)}$ would eventually result in a preference for leaving the stochastic part and going towards the already discovered goal (Condition 1 is satisfied). In other words, agents will eventually give up exploration after a sufficiently long and unsuccessful exploration phase. This is why the MB optimistic initialization is similar to exploration driven by information gain.

Moreover, by analyzing $f_{\text{C1}}$, we can gain further insights about how the model parameters influence exploration based on the MB optimistic initialization:

1. For any value of $r_2^*$ and $r_{G^*}$, we have

$$\lim_{\lambda_{\text{ext}} \to 0} f_{\text{C1}}(r_1^*, r_2^*, \lambda_{\text{ext}}, \epsilon_{\text{obs}}, \bar{C}^{(t)}, |\mathcal{S}^{(t)}|) = \infty.$$

This implies that decreasing the discount factor to put a small weight on the future rewards would make the agent stay in the stochastic part (Condition 1 is violated).

2. If $r_{G^*} < r_2^*$ (i.e., the agent knows that there exists a goal state with a reward higher than the one already discovered) and $\lambda_{\text{ext}}^2 \tilde{p}_g^{(t)} < \frac{p_s^{(t)}}{1-\lambda_{\text{ext}}(1-p_s^{(t)})}$ (i.e., the discount factor is small enough; see point 1), then we have
$$\lim_{r_2^* \to \infty} f_{\text{C1}}(r_1^*, r_2^*, \lambda_{\text{ext}}, \epsilon_{\text{obs}}, \bar{C}^{(t)}, |\mathcal{S}^{(t)}|) > r_{G^*}.$$
This implies that, if $r_{G^*} < r_2^*$, then increasing $r_2^*$ would eventually result in a preference for staying in the stochastic part (Condition 1 is violated). In other words, if the reward value of one of the three goal states is much greater than the discovered goal state, then the agent prefers to keep exploring the stochastic part.

3. For any value of $r_2^*$ and $r_{G^*}$, we have
$$\lim_{\epsilon_{\text{obs}} \to 0} f_{\text{C1}}(r_1^*, r_2^*, \lambda_{\text{ext}}, \epsilon_{\text{obs}}, \bar{C}^{(t)}, |\mathcal{S}^{(t)}|) < 0.$$
This implies that, independently of the reward value of the discovered goal state, if the agent assigns a very small prior probability to the unseen transitions, then the agent always prefer to leave the stochastic part and go to the already discovered goal state (i.e., Condition 1 is satisfied).

### 3 Algorithmic implementation

#### 3.1 Initialization

For Epi $> 1$, $\mathcal{S}^{(0)}$, $\tilde{C}^{(0)}$, $U^{(0)}_{\text{ext}}$, $U^{(0)}_{\text{int}}$, $Q^{(0)}_{\text{MB,ext}}$, $Q^{(0)}_{\text{MB,int}}$, $Q^{(0)}_{\text{MF,ext}}$, and $Q^{(0)}_{\text{MF,int}}$ are initialized by their latest value in the previous episode.

For Epi $= 1$, the initial values are as follows:

$$\begin{aligned} \mathcal{S}^{(0)} &= \{G_0, G_1, G_2\}, \\ \tilde{C}^{(0)} &= 0, \\ Q^{(0)}_{\text{MF,ext}}(s,a) &= Q^{(0)}_{\text{MF,ext}}, \\ Q^{(0)}_{\text{MF,int}}(s,a) &= Q^{(0)}_{\text{MF,int}}. \end{aligned} \tag{S32}$$

For the model-based $Q$-values, we can analytically solve the bellman equations at time $t = 0$, resulting in

$$\begin{aligned} U^{(0)}_{\text{ext}}(s) = Q^{(0)}_{\text{MB,ext}}(s,a) &= \frac{\hat{\theta}_{\text{obs}} + \lambda_{\text{ext}} \hat{\theta}_{\text{new}} W_{\text{obs}}}{1 - \lambda_{\text{ext}} |\mathcal{S}^{(0)}| \Big( \hat{\theta}_{\text{obs}} + \lambda_{\text{ext}} \hat{\theta}_{\text{new}} W_{\text{obs}} \Big)} (1 + r_1 + r_2), \\ U^{(0)}_{\text{int}}(s) = Q^{(0)}_{\text{MB,int}}(s,a) & \\ = \frac{\hat{\theta}_{\text{obs}} |\mathcal{S}^{(0)}| R^{(\text{int})}_{\text{obs}}(s_{\text{obs}}) + \hat{\theta}_{\text{new}} \Big( R^{(\text{int})}_{\text{obs}}(s_{\text{new}}) + \lambda_{\text{int}} W_{\text{new}} R^{(\text{int})}_{\text{new}}(s_{\text{new}}) + |\mathcal{S}^{(0)}| \lambda_{\text{int}} W_{\text{obs}} R^{(\text{int})}_{\text{new}}(s_{\text{obs}}) \Big)}{1 - \lambda_{\text{int}} |\mathcal{S}^{(0)}| \Big( \hat{\theta}_{\text{obs}} + \lambda_{\text{int}} \hat{\theta}_{\text{new}} W_{\text{obs}} \Big)}&, \end{aligned} \tag{S33}$$

with

$$\begin{aligned} W_{\text{obs}} &= \frac{\epsilon_{\text{obs}}}{(1-\lambda)\epsilon_{\text{new}} + \epsilon_{\text{obs}} |\mathcal{S}^{(0)}|} \quad , \quad W_{\text{new}} = \frac{\epsilon_{\text{new}}}{(1-\lambda)\epsilon_{\text{new}} + \epsilon_{\text{obs}} |\mathcal{S}^{(0)}|}, \\ \hat{\theta}_{\text{obs}} &= \frac{\epsilon_{\text{obs}}}{\epsilon_{\text{new}} + \epsilon_{\text{obs}} |\mathcal{S}^{(0)}|} \quad , \quad \hat{\theta}_{\text{new}} = \frac{\epsilon_{\text{new}}}{\epsilon_{\text{new}} + \epsilon_{\text{obs}} |\mathcal{S}^{(0)}|} \end{aligned} \tag{S34}$$

and

$$\begin{aligned} R^{(\text{int})}_{\text{new}}(s_{\text{obs}}) &= R_{\text{int},0}(s_{\text{new}}, a \to s) \quad , \quad R^{(\text{int})}_{\text{new}}(s_{\text{new}}) = R_{\text{int},0}(s_{\text{new}}, a \to s_{\text{new}}) \\ R^{(\text{int})}_{\text{obs}}(s_{\text{obs}}) &= R_{\text{int},0}(s, a \to s) \quad , \quad R^{(\text{int})}_{\text{obs}}(s_{\text{new}}) = R_{\text{int},0}(s, a \to s_{\text{new}}) \end{aligned} \tag{S35}$$

for any $a \in \mathcal{A}$ and $s \in \mathcal{S}^{(0)}$.

However, the final (after learning the transition probabilities) values for $Q_{\text{MB,ext}}(s,a)$ are much smaller than the analytic solution to the Bellman equation at $t = 0$ – due to the sparse connections and a single path to one goal state. We, therefore, use a heuristic and put $U^{(0)}_{\text{ext}}(s) = Q^{(0)}_{\text{MB,ext}}(s,a) = 0$.

#### 3.2 Pseudocode

See Algorithms 1, 2, 3, and 4 for . Note that, in all pseudocode, we use an alternative shorter notation by defining $R^{(\text{int},t)}_{s,a}(s') := R_{\text{int},t}(s, a \to s')$ and $R^{(\text{ext})}_{s,a}(s') := R_{\text{ext}}(s, a \to s')$.

**Algorithm 1** General pseudocode for algorithm

### Setting specification
Specify $$\Phi = \{\Phi^{(\text{main})}, \Phi^{(\beta)}, \Phi^{(b)}\}$$
Specify the intrinsic reward function $R_{s,a}^{(\text{int},t)}(s')$.
Specify Episode (Epi) and the set of possible actions $\mathcal{A}$.
**if** Epi $= 1$ **then**
  $\beta_{\text{MB,ext}} \leftarrow \beta^{(1)}_{\text{MB,ext}}$, $\beta_{\text{MF,ext}} \leftarrow \beta^{(1)}_{\text{MF,ext}}$, $\beta_{\text{MB,int}} \leftarrow \beta^{(1)}_{\text{MB,int}}$, and $\beta_{\text{MF,int}} \leftarrow \beta^{(1)}_{\text{MF,int}}$.
**else**
  $\beta_{\text{MB,ext}} \leftarrow \beta^{(2,r_{G*})}_{\text{MB,ext}}$, $\beta_{\text{MF,ext}} \leftarrow \beta^{(2,r_{G*})}_{\text{MF,ext}}$, $\beta_{\text{MB,int}} \leftarrow \beta^{(2,r_{G*})}_{\text{MB,int}}$, and $\beta_{\text{MF,int}} \leftarrow \beta^{(2,r_{G*})}_{\text{MF,int}}$.
**end if**
### Initialization (all variables are defined only for $s \in \mathcal{S}^{(0)}$)
Initialize $\mathcal{S}^{(0)}$, $\tilde{C}^{(0)}$, $U^{(0)}_{\text{ext}}$, $U^{(0)}_{\text{int}}$, $Q^{(0)}_{\text{MB,ext}}$, $Q^{(0)}_{\text{MB,int}}$, $Q^{(0)}_{\text{MF,ext}}$, and $Q^{(0)}_{\text{MF,int}}$ (cf. subsection 3.1).
### 1st observation
$t \leftarrow 0$
Initialize state $s_1$ and update $\tilde{C}^{(1)}_s \leftarrow \kappa \tilde{C}^{(0)}_s + \delta_{s,s_1}$ and $\mathcal{S}^{(1)} \leftarrow \mathcal{S}^{(0)} \cup \{s_1\}$.
$\tilde{C}^{(1)}_{s,a,s'} \leftarrow \tilde{C}^{(0)}_{s,a,s'}$ for all $s$ and $s' \in \mathcal{S}^{(0)}$.
$Q^{(1)}_{\text{MF,ext}}(s,a) \leftarrow Q^{(0)}_{\text{MF,ext}}(s,a)$ and $Q^{(1)}_{\text{MF,int}}(s,a) \leftarrow Q^{(0)}_{\text{MF,int}}(s,a)$ for all $s \in \mathcal{S}^{(0)}$.
$e^{(1)}_{\text{ext}} \leftarrow 0$ and $e^{(1)}_{\text{int}} \leftarrow 0$.
### Extensions of variables for $s_1 \notin \mathcal{S}^{(0)}$
$\tilde{C}^{(1)}_{s,a,s'} \leftarrow 0$ if $s = s_1$ or $s' = s_1$ and $s_1 \notin \mathcal{S}^{(0)}$.
$Q^{(1)}_{\text{MF,ext}}(s_1,a) \leftarrow Q^{(0)}_{\text{MF,ext}}$ and $Q^{(1)}_{\text{MF,int}}(s_1,a) \leftarrow Q^{(0)}_{\text{MF,int}}$ if $s_1 \notin \mathcal{S}^{(0)}$.
Update $U^{(1)}_{\text{ext}}$, $U^{(1)}_{\text{int}}$, $Q^{(1)}_{\text{MB,ext}}$, and $Q^{(1)}_{\text{MB,int}}$ using the model-based branch in Alg. 2.
### Going through the task
$t \leftarrow 1$.
**while** $s_t \neq G_i$ for $i \in \{0,1,2\}$ **do**
### Making action
  Compute $Q^{(t)}_{\text{MF}}(s,a) \leftarrow \beta_{\text{MF,ext}} Q^{(t)}_{\text{MF,ext}}(s,a) + \beta_{\text{MF,int}} Q^{(t)}_{\text{MF,int}}(s,a)$.
  Compute $Q^{(t)}_{\text{MB}}(s,a) \leftarrow \beta_{\text{MB,ext}} Q^{(t)}_{\text{MB,ext}}(s,a) + \beta_{\text{MB,int}} Q^{(t)}_{\text{MB,int}}(s,a)$.
  Sample $a_t$ with probability $\pi(a_t|s_t) \propto \exp\Big\{Q^{(t)}_{\text{MF}}(s_t,a_t) + Q^{(t)}_{\text{MB}}(s_t,a_t) + b(a_t)\Big\}$.
  Observe $s_{t+1}$.
### Updating internal variables
  $\mathcal{S}^{(t+1)} \leftarrow \mathcal{S}^{(t)} \cup \{s_{t+1}\}$.
  Update counts $\tilde{C}^{(t+1)}_s \leftarrow \kappa \tilde{C}^{(t)}_s + \delta_{s,s_{t+1}}$ and $\tilde{C}^{(t+1)}_{s_t,a_t,s'} \leftarrow \kappa \tilde{C}^{(t)}_{s_t,a_t,s'} + \delta_{s',s_{t+1}}$.
  $\tilde{C}^{(t+1)}_{s,a,s'} \leftarrow \tilde{C}^{(t)}_{s,a,s'}$ if $s \neq s_t$ or $a \neq a_t$.
  Update $U^{(t+1)}_{\text{ext}}$, $U^{(t+1)}_{\text{int}}$, $Q^{(t+1)}_{\text{MB,ext}}$, and $Q^{(t+1)}_{\text{MB,int}}$ using the model-based branch in Alg. 2.
  Update $e^{(t+1)}_{\text{ext}}$, $e^{(t+1)}_{\text{int}}$, $Q^{(t+1)}_{\text{MF,ext}}$, and $Q^{(t+1)}_{\text{MF,int}}$ using the model-free branch in Alg. 3.
### Going to the next step
  $t \leftarrow t + 1$.
**end while**

**Algorithm 2** Pseudocode for the model-based branch

$\tilde{C}_{s,a}^{(t+1)} \leftarrow \sum_{s'} \tilde{C}_{s,a,s'}^{(t+1)}$.

### Updating the world model

$\hat{\theta}_{s,a}^{(t+1)}(s') \leftarrow \big(\epsilon_{\text{obs}} + \tilde{C}_{s,a,s'}^{(t+1)}\big)/\big(\epsilon_{\text{new}} + \epsilon_{\text{obs}}|\mathcal{S}^{(t+1)}| + \tilde{C}_{s,a}^{(t+1)}\big)$ for $s' \in \mathcal{S}^{(t+1)}$.

$\hat{\theta}_{s,a}^{(t+1)}(s_{\text{new}}) \leftarrow \big(\epsilon_{\text{new}}\big)/\big(\epsilon_{\text{new}} + \epsilon_{\text{obs}}|\mathcal{S}^{(t+1)}| + \tilde{C}_{s,a}^{(t+1)}\big)$.

### Updating the values

Update $Q_{\text{MB,int}}^{(t+1)}$ and $U_{\text{int}}^{(t+1)}$ using Alg. 4 and $R^{(\text{int},t+1)}$ as rewards.

Update $Q_{\text{MB,ext}}^{(t+1)}$ and $U_{\text{ext}}^{(t+1)}$ using Alg. 4 and $R^{(\text{ext})}$ as rewards.

**Algorithm 3** Pseudocode for the model-free branch

### Prediction errors

$RPE_{\text{ext},t+1} \leftarrow R_{s_t,a_t}^{(\text{ext})}(s_{t+1}) + \lambda_{\text{ext}} \max_{a' \in \mathcal{A}} Q_{\text{MF,ext}}^{(t)}(s_{t+1}, a') - Q_{\text{MF,ext}}^{(t)}(s_t, a_t)$.

$RPE_{\text{int},t+1} \leftarrow R_{s_t,a_t}^{(\text{int},t)}(s_{t+1}) + \lambda_{\text{int}} \max_{a' \in \mathcal{A}} Q_{\text{MF,int}}^{(t)}(s_{t+1}, a') - Q_{\text{MF,int}}^{(t)}(s_t, a_t)$.

### Update of the eligibility traces

$e_{\text{ext}}^{(t+1)}(s_t, a_t) \leftarrow 1$, and $e_{\text{ext}}^{(t+1)}(s, a) \leftarrow \lambda_{\text{ext}}\mu_{\text{ext}} e_{\text{ext}}^{(t)}(s, a)$, for all $s \neq s_t$ and $a \neq a_t$.

$e_{\text{int}}^{(t+1)}(s_t, a_t) \leftarrow 1$, and $e_{\text{int}}^{(t+1)}(s, a) \leftarrow \lambda_{\text{int}}\mu_{\text{int}} e_{\text{int}}^{(t)}(s, a)$, for all $s \neq s_t$ and $a \neq a_t$.

### TD-learners

$Q_{\text{MF,ext}}^{(t+1)}(s, a) \leftarrow Q_{\text{MF,ext}}^{(t)}(s, a) + \rho e_{\text{ext}}^{(t+1)}(s, a) RPE_{\text{ext},t+1}$, $\forall s \in \mathcal{S}$ and $a \in \mathcal{A}$.

$Q_{\text{MF,int}}^{(t+1)}(s, a) \leftarrow Q_{\text{MF,int}}^{(t)}(s, a) + \rho e_{\text{int}}^{(t+1)}(s, a) RPE_{\text{int},t+1}$, $\forall s \in \mathcal{S}$ and $a \in \mathcal{A}$.

**Algorithm 4** Pseudocode for the modified Prioritized Sweeping Algorithm at time $t+1$

### Specifying whether the update is for the intrinsic or the extrinsic reward

$\lambda \leftarrow \lambda_{\text{ext}}$ for extrinsic and $\lambda \leftarrow \lambda_{\text{int}}$ for intrinsic reward.

$Q^{(t)} \leftarrow Q^{(t)}_{\text{MB,ext}}$, $U^{(t)} \leftarrow U^{(t)}_{\text{ext}}$, and $R \leftarrow R^{(\text{ext})}$ for extrinsic, and $Q^{(t)} \leftarrow Q^{(t)}_{\text{MB,int}}$, $U^{(t)} \leftarrow U^{(t)}_{\text{int}}$, and $R \leftarrow R^{(\text{int},t+1)}$ for intrinsic reward.

### Extending $U$-values

$W_{\text{obs}} \leftarrow \epsilon_{\text{obs}} / \big((1-\lambda)\epsilon_{\text{new}} + \epsilon_{\text{obs}}|\mathcal{S}^{(t+1)}|\big)$ and $W_{\text{new}} \leftarrow \epsilon_{\text{new}} / \big((1-\lambda)\epsilon_{\text{new}} + \epsilon_{\text{obs}}|\mathcal{S}^{(t+1)}|\big)$

**if** $s_{t+1} \notin \mathcal{S}^{(t)}$ **then**

    $U^{(t)}(s_{t+1}) \leftarrow W_{\text{obs}} \sum_{s' \in \mathcal{S}^{(t)}} \Big(R_{s_{\text{new}}}(s') + \lambda U^{(t)}(s')\Big) + W_{\text{new}} R_{s_{\text{new}}}(s_{\text{new}})$.

**end if**

### Applying the effect of the latest observation on $Q$-values using previous $U$-values

**for** $(s,a) \in \mathcal{S}^{(t+1)} \times \mathcal{A}$ **do**

    $Q^{(t+1)}(s,a) \leftarrow Q\text{Update}(s,a;\lambda,\hat{\theta}^{(t+1)},U^{(t)},R,W_{\text{obs}},W_{\text{new}},\mathcal{S}^{(t+1)})$ defined in Eq. S16.

**end for**

### Making the priority queue

**for** $s \in \mathcal{S}^{(t+1)}$ **do**

    $U^{(t+1)}(s) \leftarrow U^{(t)}(s)$

    $\text{PriorityQueue}(s) \leftarrow |U^{(t+1)}(s) - \max_{a\in\mathcal{A}} Q^{(t+1)}(s,a)|$

**end for**

### Updating $U$-values for $T_{\text{PS}}$ steps

**for** $T_{\text{PS}}$ iterations **do**

    $s' \leftarrow \arg\max_{s\in\mathcal{S}^{(t+1)}} \text{PriorityQueue}(s)$

    $\Delta V \leftarrow \max_{a\in\mathcal{A}} Q^{(t+1)}(s',a) - U^{(t+1)}(s')$

    $U^{(t+1)}(s') \leftarrow \max_{a\in\mathcal{A}} Q^{(t+1)}(s',a)$

    # Applying the effect of the update of $U$-values on $Q$-values

    **for** $(s,a) \in \mathcal{S}^{(t+1)} \times \mathcal{A}$ **do**

        $Q^{(t+1)}(s,a) \leftarrow Q^{(t+1)}(s,a) + \lambda\Big(\hat{\theta}^{(t+1)}_{s,a}(s') + \lambda\hat{\theta}^{(t+1)}_{s,a}(s_{\text{new}})W_{\text{obs}}\Big)\Delta V$

    **end for**

    # Updating the priority queue

    **for** $s \in \mathcal{S}^{(t+1)}$ **do**

        $\text{PriorityQueue}(s) \leftarrow |U^{(t+1)}(s) - \max_{a\in\mathcal{A}} Q^{(t+1)}(s,a)|$

    **end for**

**end for**

Schuurmans, D., Bengio, Y., and Bottou, L., editors, *Advances in Neural Information Processing Systems*, volume 21. Curran Associates, Inc.
